# Functional dissection of the conserved *C. elegans* LEM-3/ANKLE1 nuclease reveals a crucial requirement for the LEM-like and GIY-YIG domains for DNA bridges processing

**DOI:** 10.1101/2024.06.26.600907

**Authors:** Junfang Song, Peter Geary, Ye Hong, Stéphane G.M. Rolland, Anton Garter

## Abstract

Faithful chromosome segregation requires the removal of all DNA bridges physically linking chromatids before the completion of cell division. While several redundant safeguard mechanisms to process these DNA bridges exist from S-phase to late anaphase, the conserved LEM-3/ANKLE1 nuclease has been proposed to be part of a ‘last chance’ mechanism that acts at the midbody to eliminate DNA bridges that persist until late cytokinesis. We show that LEM-3 can cleave a wide range of branched DNA substrates, including flaps, forks, nicked and intact Holliday Junctions. AlphaFold modeling data suggest that the catalytic mechanism of LEM-3/ANKLE1 is conserved, mirroring the mechanism observed in bacterial GIY-YIG nucleases. We also present evidence that LEM-3 may form a homodimeric complex on the Holliday Junction DNA. LEM-3 DNA binding capacity requires both the LEM-like and the GIY-YIG nuclease domains; both are also essential for LEM-3 recruitment to the midbody and its nuclease activity. Finally, we show that preventing LEM-3 nuclear access is important to avoid toxicity, likely caused by branched DNAs cleavage during normal DNA metabolism. Our data suggest that *C. elegans* LEM-3 acts as a ‘last chance catch-all’ enzyme that processes DNA bridges caused by various perturbations of DNA metabolism just before cells divide.

## Introduction

Faithful cell division requires the removal of all DNA connections that link segregating chromatids. Such connections result from branched DNA species that arise from perturbations in DNA metabolism, including DNA under-replication, persistent intermediates of recombinational repair, or DNA entanglement caused by topoisomerase deficiency or compromised chromosome condensation (1). DNA bridges also occur when bicentric chromosomes, arising from telomere-crisis or reciprocal translocations, are pulled between opposite poles, a process initially observed by Barbara McClintock (2). During mitosis, DNA connections cytologically manifest as DNA bridges. Chromatinized bridges are referred to as chromatin bridges. When not packaged into chromatin, DNA bridges are called ultrafine bridges (UFBs) (3). Bridges are commonly observed during anaphase but can persist into late telophase. Bridges can extensively delay the completion of cytokinesis by triggering the cytokinesis checkpoint. Defective bridge resolution can lead to cytokinesis failure, tetraploidization, and aneuploidy (4–7). When persistent bridges rupture during cytokinesis, micronuclei formation, often associated with chromothripsis, the localized scattering of chromosomes followed by random assembly of fragments, occurs (8, 9). Therefore, the resolution of chromatin bridges is of pivotal importance for genomic integrity.

4-way branched DNA structures referred to as Holliday Junctions (HJs) are central intermediates of recombinational repair, resolved by multiple redundant mechanisms. During S-phase, a complex that includes the Blooms helicase and topoisomerase I dissolves adjoining HJs. In G2, persistent junctions are processed by the SLX4 scaffold protein in association with the SLX1 and the EME1/MUS81 nucleases. SLX1 catalyzes a nick, the nicked HJ being the preferred substrate for MUS81 (10). In anaphase, the remaining junctions are resolved by the GEN1 resolvase (11, 12). Despite these redundant mechanisms, some bridges persist beyond anaphase, and recent evidence suggests that the conserved *C. elegans* LEM-3 nuclease and its human ortholog ANKLE1 process these remnant bridges (13, 14).

Maintaining genome stability in the first zygotic cell divisions is exceptionally challenging, particularly so in the *C. elegans* nematode, where embryonic cell cycles occur in rapid succession, with the S-phase preceding the first zygotic cell division requiring only ∼8.5 minutes (15, 16). Analysis of the *C. elegans* LEM-3 nuclease indicated that LEM-3 acts as part of a ‘last-chance’ mechanism to process DNA bridges right before cells divide (13). LEM-3 dynamically localizes to the midbody, the structure where cells abscise just before the completion of cytokinesis. The early stages of cytokinesis, which involves the assembly of the central spindle and midbody formation, together with LEM-3 midbody localization, are required for DNA bridge processing to maintain genome stability. Real-time imaging of *C. elegans* embryos showed that chromatin bridges caused by persistent recombination intermediates, DNA under-replication, or chromosome entanglement caused by partial condensin depletion persist in *lem-3* mutant embryos while they are processed in wild-type embryos (13). These results and the exquisite sensitivity of *lem-3* mutants to multiple DNA-damaging agents indicate that LEM-3 might have a broad substrate specificity to contribute to genome stability. The substrate specificity of LEM-3 has yet to be determined.

Knockout mice of *Ankle1* (the vertebrate LEM-3 ortholog) do not have an overt phenotype, and cell lines derived from these mice do not show hypersensitivity towards various DNA-damaging agents (17). More recently, ANKLE1 has been shown to localize to the midbody like its nematode ortholog (14). Furthermore, ANKLE1 defective cancer cell lines have been shown to be moderately hypersensitive to treatment with chemotherapeutic drugs, including cisplatin, a DNA cross-linking agent, and ICRF-193, a topoisomerase II inhibitor, that traps two DNA double strands within the topoisomerase complex (14). ANKLE1 has been shown to have a structure-specific endonuclease activity (14, 18–20) and, at high concentrations, to lead to exonucleolytic degradation (14, 18–20). The enzyme also cleaves a broad range of branched DNA substrates *in vitro*, including splayed DNA structures and HJ (20). To cleave DNA bridges, ANKLE1 was proposed to nick dsDNA, which would provide access to the TREX1 exonuclease for ssDNA generation, with TREX activity on opposing strands leading to the breakage of a double-stranded bridge (14). Such a mode of bridge cleavage would be error-prone, in line with the evidence (derived from clones that survive bridge breakage) for chromothripsis and kataegis, the latter process characterized by locally-clustered APOBEC deaminase activity on ssDNA substrates (21). Nevertheless, ANKLE1 activity has been suggested to prevent even more deleterious chromosome fragmentation events, which occur when DNA ruptures during cell division (14). All in all, vertebrate evolution might have prioritized ANKLE1 bridge cleavage to allow for cytokinesis, even if it is associated with a risk of chromosomal aberrations. In addition, ANKLE1 has been reported to function in mitochondrial DNA degradation, which occurs during erythrocyte differentiation (22). In contrast, the exquisite sensitivity of *lem-3* mutant worms to various forms of DNA damage and the requirement for LEM-3 midbody localization for mending branched DNA intermediates suggest that *C. elegans* LEM-3 may be part of a mechanism that promotes genome integrity (9).

LEM-3/ANKLE1 and SLX-1 are the only Eukaryotic GIY-YIG nucleases, with LEM-3/ANKLE1 being encoded only in animals. LEM-3 is composed of 704 amino acids and, like ANKLE1, contains three distinct domains (Figure 1A): an N-terminal Ankyrin repeat domain, a central LEM domain, and a C-terminal ∼100 amino-acid GIY-YIG nuclease domain, a domain that features a conserved core (23, 24). GIY-YIG nuclease domains consisting of three central β-sheet strands and two helices are shared by prokaryotic homing endonucleases (such as I-TevI and I-BmoI), which are encoded within introns or inteins of host genes and can specifically recognize and cleave target DNA sequences (25)(23). Other members, such as T4 EndoII (26) and REases, remove foreign DNA by cleaving specific target sites (27). The bacterial UvrC endonuclease participates in the nucleotide excision repair pathway by incising the damaged DNA strand (28).

**Figure 1:**
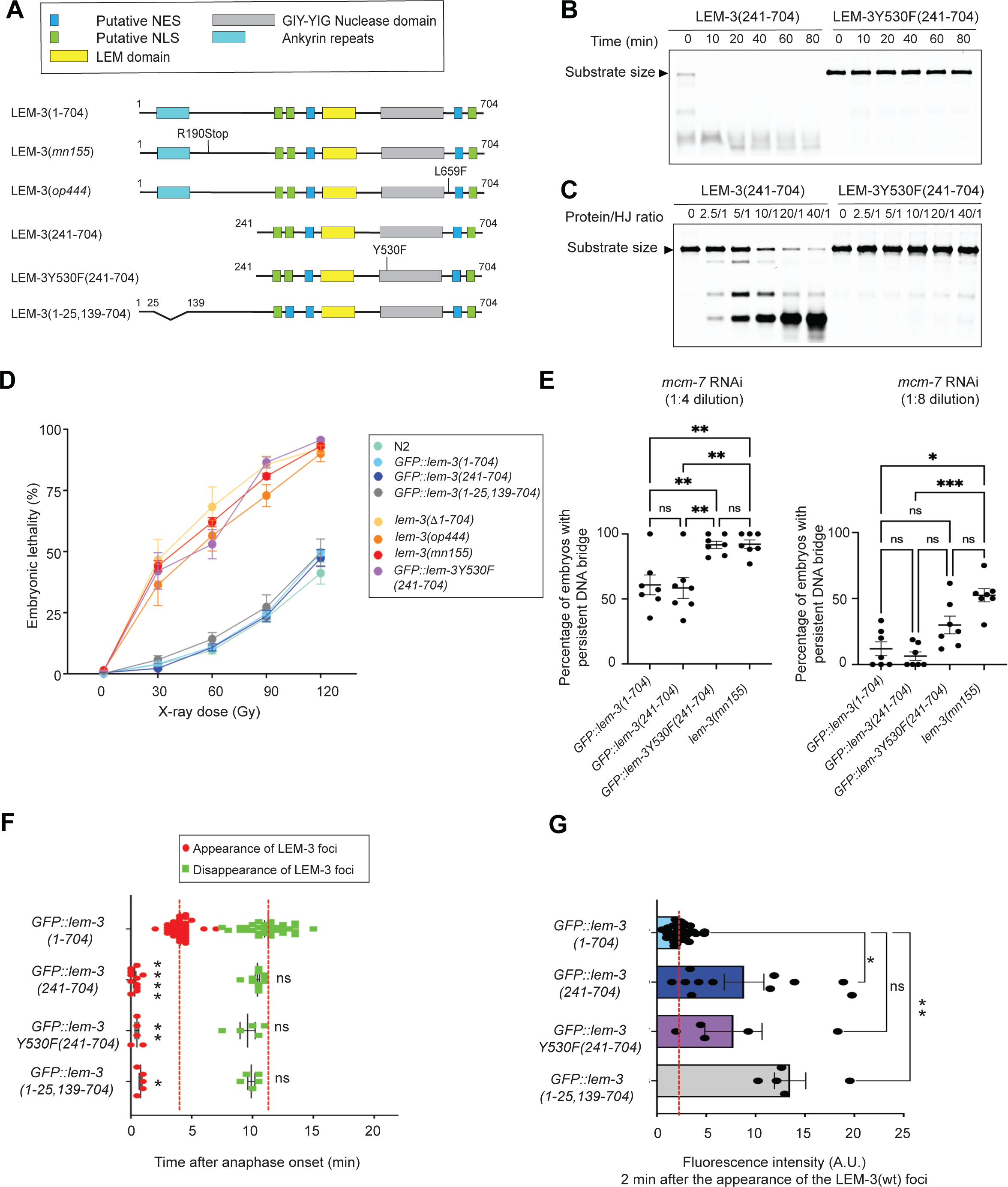
*in vitro* and *in vivo* characterization of LEM-3(241-704). **(A)** Schematic of the different LEM-3 derivatives used in this figure. **(B-C)** Dose- and time-dependent cleavage of HJ substrate by LEM-3(241-704) or LEM-3Y530F(241-704). In panel A, a protein/HJ molar ratio of 50/1 was used. In panel B, incubation at 37℃ was performed for 10 mins. The cleavage products were analyzed by 6% neutral PAGE. The size of the DNA substrate is indicated. **(D)** Analysis of the IR sensitivity of the indicated LEM-3 derivates (n≽5, mean and SEM). **(E)** Analysis of the sensitivity to partial *mcm-7* RNAi of the indicated LEM-3 derivates. Different dilutions of RNAi were used (*mcm-7* RNAi was diluted 1:4 and 1:8 with *mock* RNAi) and the percentage of 2-cell stage embryos with persistent DNA bridges was determined (n=7 experiments (in each experiment ∼10-15 embryos were analyzed), mean and SEM are shown. For 1:4 dilution, ns = not significant and ** = p <0.01 by one-way ANOVA with Tukey’s multiple comparisons test. For 1:8 dilution, ns = not significant and * p < 0.05 and *** p < 0.001 by Kruskal-Wallis with Dunn’s multiple comparison). **(F)** Measurement of the time of appearance and disappearance of the LEM-3 foci for the indicated LEM-3 derivates (n≽5, mean and SEM. For the time of appearance, * p < 0.05, ** p < 0.01, and **** p < 0.0001 by Kruskal-Wallis with Dunn’s multiple comparison to *GFP::lem-3(wt)*. For the time of disappearance, ns = not significant by Kruskal-Wallis with Dunn’s multiple comparisons to *GFP::lem-3(wt)*). **(G)** The fluorescence intensity of the indicated LEM-3 derivates foci was measured 2 minutes after the appearance of the wild-type LEM-3 foci (n≽5, mean and SEM. ns = not significant, * p < 0.05 and ** p < 0.01 by Brown-Forsythe and Welch ANOVA tests with Dunnett’s T3 multiple comparisons to *GFP::lem-3(wt)*).

Nucleases involved in DNA repair need to be spatially and temporally controlled, in order not to cleave DNA structures inappropriately. LEM-3, like ANKLE1, is excluded from the nucleus, and ANKLE1 forced nuclear expression leads to genome instability (18). However, how the activity and localization of LEM-3/ANKLE1 is regulated remains largely unclear. Here, taking advantage of the *C. elegans* experimental system, with *lem-3* mutations leading to exquisite DNA damage sensitivity and real-time imaging facilitated by rapid cell cycle progression, we report on an *in vivo* structure-function analysis of the *C. elegans* LEM-3 nuclease. We also characterize substrate specificity and basic reaction mechanisms.

### Materials and Methods Experimental model

*Caenorhabditis elegans (C. elegans)* strains were maintained at 20°C unless stated otherwise for experimental purposes. They were kept on nematode growth medium (NGM) plates that were seeded with OP50 bacteria (100μl per medium plate). The list of strains used in this study is provided in Table S1.

### CRISPR-Cas9 Genome Edits

As indicated in Table S1, some genome edits were generated by Sunybiotech, and some were generated in the Gartner lab following the previously published Mello and co-workers methodology (57). For the generation of each genome edit, ∼10-30 young adult hermaphrodites were injected in one gonad arm. They were then recovered individually into 5μl of M9 buffer in the center of the OP50 seeded plate. Recovered worms were then maintained at 20°C. F1 progeny that displayed the roller phenotype were singled 3-4 days post-injection and subsequently screened by using PCR and/or sequencing after being permitted to lay eggs for 24-48 hours. For the edits generated in the Gartner lab, the crRNA and ssODN used are indicated in Table S2.

### L4 survival assay

Experiments were conducted in triplicate. L1 filtration was conducted from freshly starved plates in order to obtain a synchronized population, as previously described (58). This was achieved by washing each medium NGM plate with 2ml of M9. A 20ml syringe was used to recover the non-synchronized population of worms from the plate. The solution was then filtered through a nylon mesh, with the size of the holes only permitting L1 larvae to pass through. The L1 filtration was done ∼72 hours prior to the IR treatment, with the *lem-3(mn155)* and *lem-3(485-704)* strains being the exception, filtered ∼84 hours prior. Following the filtration, the L1 larvae were transferred in 2 separate drops of 100μl onto NGM plates. After allowing the plates to dry, they were incubated at 15°C until the worms reached the L4 larvae stage. L4 larvae were treated with different IR doses (0Gy, 30Gy, 60Gy, 90Gy, 120Gy) using a Biological X-ray irradiator (Rad Source - RS 2000). After IR, the worms were allowed to recover at 20°C for 24 hours. 5 adult hermaphrodites for every strain (except for the *lem-3(mn155)* and *lem-3(485-704)* strains where 10 adult hermaphrodites were used) were then transferred to seeded medium NGM plates. They were allowed to lay eggs for 6 hours at 20°C and then removed from the plates. Subsequently, the number of eggs laid on each plate was counted. The embryos were then incubated at 20°C for 24 hours before counting the unhatched (dead) eggs. The following day, the number of unhatched eggs was recounted, as was the number of alive larvae. The embryonic survival was scored 72 hours after IR as the ratio of dead eggs to total laid eggs.

### Confocal Image Acquisition

Embryos from 4-5 adult hermaphrodites were dissected in M9 and mounted on 2% agarose pads, similar to what has been done previously (13). Of note, we used for all experiments, high precision 18x18 coverslip (1.5H Zeiss). Z stacks (15 slices; 1.5μm per step) were captured every 30 seconds for 30 minutes at room temperature (RT) (20°C to 25°C) using a spinning-disk confocal microscope (ECLIPSE Ti2-E, Nikon) with spinning disk head (CSU-W1, Yokogawa Electric Corporation) and the NIS element software (Nikon). A 60x oil objective (NA 1.4) was used to capture the images. Embryos were sequentially illuminated by light from 488nm and 561nm laser, with an exposure time of 300ms, and the power of each laser was adjusted to 50% of its maximal capacity. 2 times line averaging was used during the imaging. Image analysis and video processing were performed using Fiji software, with the signal analyzed both 2 mins post signal onset and at the same time point measured in the control, provided the signal of our mutant was already present at the control signal onset. The time of onset is relative to anaphase onset of the respective embryo, and a minimum of 5 embryos were analyzed for each strain in each experimental condition.

### RNAi

Ahringer RNAi clone for *mcm-7* was obtained from Source Bioscience. The RNAi bacteria containing the empty L4440 vector was used as mock RNAi. In order to prepare the RNAi plates, *mcm-7* and mock RNAi bacteria were inoculated in LBC (LB supplemented with 100μg/ml of Carbenicillin) and grown overnight at 37°C. Each culture was then diluted to obtain the desired OD_600_ of 0.5, after which 50μl was used to seed the RNAi plates (NGM plates supplemented with 6mM IPTG and 100μg/ml of Carbenicillin). The RNAi plates were then placed under aluminum foil as IPTG is light-sensitive and incubated under the hood overnight. The next day, ∼30 P0 L4 larvae were transferred onto the RNAi plates and incubated at 25°C. 24 hours later, F1 embryos were analyzed by confocal microscopy as described in the ‘confocal Image Acquisition’ section above.

### Partial RNAi experiment

As with the RNAi protocol, RNAi cultures were grown in LBC overnight at 37°C, with a final OD_600_ of 0.5 used to seed the RNAi plates. Prior to analysis, L4 larvae (∼25-30) were transferred onto RNAi plates and incubated at 25°C for 24 hours. At this point, the RNAi plates were re-labelled, so the analysis could be conducted blind. The next day, microscope slides were cleaned with 70% ethanol and labeled with the corresponding new labels of the RNAi plates. Following this, each microscope slide was coated twice with poly-L-lysine (0.25mg/ml), and dried in between coats on a heat plate at 80°C. After the second coat was added, the slides were given appropriate time to cool down before the next step. Once the slides had cooled down, 10μL of ultrapure water (Sigma) was added to the center of an 18x18 coverslip (1.5H Zeiss). Adult P0 hermaphrodites from the corresponding RNAi-labeled plate were then transferred into the drop of water. They were then cut open, allowing the F1 embryos to be released. The embryos were then transferred using a P10 pipette to the coated microscope slide. A new 18x18 coverslip (1.5H Zeiss) was then placed on top of the area where the embryos were transferred (at ∼45° angle, with the corner of the coverslip hanging over the edge of the coated microscope slide). The microscope slide was then immediately flash-frozen on dry ice (for at least 30 mins). The coverslip was then very quickly removed by flicking it (freeze-crack method), and the slides were incubated in methanol/acetone (1:1) for ∼10 mins. Following the fixation, the slides were air-dried (for ∼8 mins) before being stored at -20°C if the next step was not performed right away. Prior to staining with Vectashield (which contains DAPI), the slides were washed in PBST for ∼10 mins. With the exception of the area containing the embryo, the rest of the slide was dried using a KIMTECH tissue. 10μL of Vectashield was then added to the area containing the embryo, and an 18x18 coverslip (1.5H Zeiss) was placed on top. Nail polish was then used to seal the slides, with the nail polish allowed to dry in the dark (as Vectashield + DAPI is light sensitive). Once all slides were sealed correctly, they were then placed at 4°C until analysis.

Slides were analyzed using a Zeiss Imager M2 microscope equipped with a 63x oil objective (NA 1.25) and a 465nm LED (for DAPI). ∼10-15 2-cell stage embryos were analyzed per slide and the percentage of 2-cell stage embryos with persistent DNA bridges was determined.

### Immunostaining of *C. elegans* embryos

Microscope slides were cleaned with 70% ethanol. Following this, each microscope slide was coated twice with poly-L-lysine (0.25mg/ml), and dried in-between coats on a heat plate at 80°C. After the second coat was added, the slides were given appropriate time to cool down before the next step. Once the slides had cooled down, 10μL of ultrapure water (Sigma) was added to the center of an 18x18 coverslip (1.5H Zeiss). Adult hermaphrodites (24 hours post L4) were then transferred into the drop of water. They were then cut open, allowing the embryos to be released. The embryos were then transferred using a P10 pipette to the coated microscope slide. A new 18x18 coverslip (1.5H Zeiss) was then placed on top of the area where the embryos were transferred (at ∼45° angle, with the corner of the coverslip hanging over the edge of the coated microscope slide). The microscope slide was then immediately flash-frozen on dry ice (for at least 30 mins). The coverslip was then very quickly removed by flicking it (freeze-crack method), and the slides were incubated in methanol/acetone (1:1) for ∼10 mins. Following the fixation, the slides were air-dried (for ∼8 mins) before being stored at -20°C if the next step was not performed right away. Permeabilization of the embryos was performed by incubating the slides 4 times 10 mins with PBS Triton 1%. After 3 washes with PBS for 10 mins, the slides were blocked with PBSTB (PBS + tween 0.1% BSA 1%) for 30 mins at RT. Primary antibody (anti-GFP (Roche) 1:500) was added to the embryos, and incubation was performed overnight at 4°C in a humid chamber. After 3 washes with PBST (PBS + tween 0.1%) for 10 mins, a secondary antibody (Alexa 488 anti-mouse (Invitrogen) 1:500) was added to the embryos, and incubation was performed for 2 hours at RT in a humid chamber. After 2 washes with PBST and 1 wash with PBS, post-fixation with PBS + 3.7% PFA was performed for 10 mins. The slide was then washed with PBS for 10 mins, with PBST for 2 times 10 mins, and with PBS for 10 mins. Afterward, with the exception of the area containing the embryo, the rest of the slide was dried using a KIMTECH tissue. 10μL of Vectashield was then added to the area containing the embryo, and an 18x18 coverslip (1.5H Zeiss) was placed on top. Nail polish was then used to seal the slides, with the nail polish allowed to dry in the dark. Once all slides were sealed correctly, they were then placed at 4°C until analysis.

For image acquisition, embryos were analyzed using a Zeiss Imager M2 microscope. A 63x oil objective (NA 1.25) was used to capture the images, with a Z stack with 0.5μm per step used. Embryos were sequentially illuminated by 465nm LED (for DAPI) and 488nm LED (for Alexa 488 anti-mouse), with an exposure time of 200ms, and the power of the LED for DAPI and Alexa 488 was adjusted to 5% or 20% of its maximal capacity, respectively. Images were captured with an Axiocam 503 camera and the software ZEN black (Zeiss). Images were deconvolved with the software ZEN black.

### Western blot analysis of LEM-3::3xHA tagged proteins from *C. elegans* extracts

Experiments were conducted in duplicate. 15 L4 larvae of the different strains were inoculated onto large NGM plates and incubated at 25°C for 3 days. The progenies were washed off the plates with 5 ml M9 buffer and transferred to a 15 ml Falcon tube. The animals were let to settle down for 5 min at RT. The supernatant was discarded, and the pellet was washed with 5 ml of M9 buffer. The animals were let to settle down for 5 min at RT. The supernatant was discarded, and the pellet was resuspended in 1 ml of M9 buffer and transferred to a 1.5 ml Eppendorf tube. The animals were let to settle down for 5 min at RT. The supernatant was discarded, and 2 volumes of M9 buffer and 1 volume of Laemmli buffer 4x were added. The tubes were incubated at -80°C for at least 10 min and incubated at 95°C for 5 mins. After centrifugation at 13,000 rpm for 1 min, 10μl of each sample was loaded on a 7.5% polyacrylamide SDS-PAGE gel. The gel was run at 75V for 2-3 hours. Wet transfer onto PVDF membranes was performed at 4°C at 25V (constant) for 16 hours. The membrane was blocked with PBS + 0.1% tween + 5% Milk for 1 hour at RT. Monoclonal anti-tubulin (1:2000; DM1A) and monoclonal anti-HA (1:2000; 16B12) antibodies were used as primary antibodies. As secondary antibodies, horseradish peroxidase-conjugated goat anti-mouse antibodies (BioRad #1706516) were used at 1:10000. Western was developed using ECL SuperSignal West Femto (ThermoScientific), and images were acquired using the ChemiDoc XRS+ System (Bio-Rad).

### Protein expression and purification

The LEM-3 open reading frame (ORF) codon-optimized for baculovirus expression was synthesized by GeneArt Gene Synthesis (Thermofisher), and TA cloned into pGEM-T Easy (Promega, Supplementary Information). A cDNA sequence (flanked by BamH1 and EcoRI) encoding for a His-GST tag and a TEV cleavage site was appended to the 5’ end of LEM-3. The verified His-GST-TEV-LEM-3^1-704^ DNA sequence was subcloned into MultiBac expression vector pFL using BamHI and HindIII. Expression vectors for truncated LEM-3 proteins (LEM-3^241-704^, LEM-3^341-704^, LEM-3^485-704^), as well as point mutants (LEM-3^241-704^ - Y530F, LEM-3^241-704^-R456A, LEM-3^241-704^-K437A-R456A) were generated using a Q5 Site-directed mutagenesis Kit, NEB (Table S3). For bacmid generation and protein purification, we followed standard protocols (59). His-GST-TEV-LEM-3 plasmids were transformed into DH10EMBacY, for integration into bacmids. Freshly purified bacmids were transfected into insect sf9 cells at a density of 0.6 x 10^6^ cells/ml, using the X-tremeGENE^TM^ HP DNA Transfection Reagent (Roche). Initial virus (V0) preparations were harvested 60 hours post-transfection. Production virus preparations (V1 and V2) were prepared by infecting insect cell cultures with V0 and V1, respectively. For protein expression, sf9 or sf21 cells (3L at a cell density of 1 x 10^6^/ml) were infected with baculovirus V2 at a ratio of 1:4000 to 1:1000. At day 3 of proliferation arrest (Dpa3) cells were harvested. Cell pellets were resuspended in a high salt lysis buffer consisting of 50mM Tris-Cl, pH7.5, 500mM NaCl, 1mM Tris(2-carboxyethyl) phosphine (TCEP, Sigma), EDTA-free Protease Inhibitor Cocktail (cOmplete™, Roche). The suspensions (20-30ml) were incubated on ice for 10 mins, dounced with Pestle B for 20 strokes (Dounce Homogenizer, VWR), and incubated for another 10 mins on ice before centrifugation at 30,000g for 30 mins at 4°C to remove insoluble material.

The soluble extracts (150 ml/3L culture) were gently mixed with 3ml of Glutathione Sepharose® 4B (Cytiva) pre-washed thoroughly with 10 times resin volume of ddH2O and lysis buffer, and incubated for 1h at 4°C. GST resins were subsequently washed with a wash buffer (50mM Tris-Cl, pH7.5, 500mM NaCl, 1mM TCEP). After the wash, the resins were incubated with 1/20-1/50 volume of TEV-GST protease (NEB) overnight at 4°C to remove the His-GST tag. The next day, untagged LEM-3 was eluted in wash buffer, concentrated (Protein Concentrators, ThermoFisher), and loaded onto a HiLoad 16/600 Superdex 200 pg size fractionation column (Cytiva) using an ÄKTAprime plus chromatography system (Cytiva). LEM-3 proteins were eluted in 1.2ml fractions with 120ml elution buffer (50mM Tris-Cl, pH 7.5, 500mM NaCl, 1mM TCEP, 5% glycerol). Peak fractions were examined by SDS-PAGE, and gel bands were cut out for protein identification using mass spectrometry. The fractions containing LEM-3 were pooled, concentrated, and adjusted to a final glycerol concentration of 50%. Protein concentrations were determined using the Bradford assay (BioRad). Purified proteins were aliquoted and stored at -20℃.

### DNA substrate preparation and nuclease assays

The oligonucleotides for DNA substrate preparation were synthesized and purified by Sigma UK. The oligonucleotides with 5’-Cy5-end-label were purified by PAGE, while the unlabelled ones were purified by HPLC. The sequences of oligonucleotides are listed in Table S4.

To prepare branched and linear DNAs, the Cy5-labeled oligonucleotide with a final concentration of 830nM was combined with a threefold excess of each of the other oligonucleotides (2500nM each) in annealing buffer (10mM Tris-Cl(pH7.5), 50mM NaCl in a volume of 120ul). The mixture was incubated in a water bath at 95°C for 5 mins and then allowed to cool down slowly overnight to RT by switching off the water bath.

To purify branched DNAs, DNA substrates were mixed with native 5 x native gel loading buffer (10mM Tris-Cl (pH 7.5), 50mM NaCl, and 50% glycerol). The mixture was then loaded to 6% neutral acrylamide/bis-acrylamide (49:1, Sigma A0924) gels using a Bio-Rad protean II xi cell system (30cm gel length, 1.5mm gel thickness). The gels were run for 5 hours at 200V in a cold room in 0.2 x TBE buffer. After finishing the run, the DNA substrate bands were detected using a Typhoon scanner (GE) excised and ground using a new blade, and eluted in an annealing buffer with gentle shaking overnight. The concentrations of eluted DNA substrates were measured using a NanoDrop® ND-1000 UV-Vis Spectrophotometer (ThermoFisher). Branched DNA substrates (830 nM) were stored in aliquots at -20°C.

Nuclease assays were carried out in a reaction volume of 20ul by incubating the indicated amounts of LEM-3 protein and 8.3nM 5’-Cy5-labeled DNA substrate (diluted from 830nM stock) in reaction buffer (10mM Tris-HCl, 100mg/ml BSA, 1mM MnCl_2_, 100mM NaCl) at 37℃. The reactions were stopped by adding EDTA and proteinase K to the final concentrations 1mM and 0.5mg/ml, respectively. The cleavage products were deproteinized 2 hours at RT and subsequently separated by 6% neutral polyacrylamide gel electrophoresis (PAGE) in 0.5 x TBE buffer The gels were imaged and analyzed using a Typhoon FLA9000 scanner and ImageQuant software from GE Healthcare.

For the cruciform assay, cruciform plasmid pHRX3 (provided by David Lilley) was prepared in supercoiled form using a Qiagen Plasmid Midi Kit, and then aliquoted and stored in dry form at -80℃ (60). The cruciform cleavage reactions were performed by incubating 1nM cruciform plasmid and the indicated amounts of LEM-3 protein in the reaction buffer at 37℃. The cleavage products were separated in 1.0% agarose gels in 1 x TBE buffer. The gels were stained with SYBR™ Gold (Invitrogen) and imaged by a Typhoon scanner.

### Electrophoretic mobility shift assay

8.3 nM HJ-X0-1 was mixed with the indicated concentrations of LEM-3 protein in binding buffer (10mM Tris-HCl, 100mM NaCl, 5% glycerol, and 5ng/ml poly[dI-dC]) in a volume of 10 or 20ul. After 1 hour incubation at 4°C, the samples were immediately loaded onto a 6% neutral PAGE in 0.5x TBE buffer in a cold room. The gels were run at 4°C and then imaged using a Typhoon scanner.

### AlphaFold model prediction and structure analysis

The structure of LEM-3 was predicted using ColabFold v1.5.2: AlphaFold2 (https://colab.research.google.com/github/sokrypton/ColabFold/blob/main/AlphaFold2.ipynb) (61), and AlphaFold3 (62). The predicted model with the highest prediction confidence score is shown in the main figure. The AlphaFold model was analyzed and compared to other known protein structures using PyMOL and Chimera.

### Statistical analysis

Statistical analysis was performed with Prism 9 (GraphPad Software, LLC). The normal distribution of the data was tested by a Shapiro-Wilk test. The equal variance was tested using a Brown-Forsythe test. Data showing a normal distribution and equal variance was analyzed using a one-way ANOVA with Tukey’s multiple comparison test. Data showing a normal distribution but not an equal variance was analyzed using the Brown-Forsythe ANOVA test with Dunnett’s T3 multiple comparisons. Data without normal distribution was analyzed using a Kruskal-Wallis with Dunn’s multiple comparison test.

## Results

### Purification of LEM-3(241-704) and genetic characterization

To assay LEM-3 activity *in vitro*, we aimed to purify full-length wild-type LEM-3 using baculovirus-based expression system in insect cells. While we encountered difficulty producing soluble full-length LEM-3 protein in the desired yield and quality, we were able to robustly express and purify a LEM-3 fragment comprised of aa241-704, which includes both the LEM and GIY-YIG domains (Figure S1A). Testing LEM-3(241-704) activity against a synthetic HJ substrate showed dose- and time-dependent cleavage of this substrate, and cleavage was largely alleviated when LEM-3Y530F(241-704), carrying the Y530F catalytic site mutation, was used (for further explanation see below) (Figure S1B, Figure 1A-C). Before further characterizing nuclease specificity and reaction mechanisms, we wished to confirm that LEM-3(241-704) is functional *in vivo* and, like the full-length protein, associated with the midbody during cytokinesis. We had previously generated a *GFP*::*lem-3(1-704)* fusion that fully complements the wild-type locus and deleted sequences coding for aa2-240 by genome editing to generate *GFP::lem-3(241-704).* These nematodes were protected from ionizing irradiation (IR), equally to those carrying wild-type *GFP::lem-3(1-704),* as revealed by treating hermaphrodite worms with IR at the L4 developmental stage and scoring embryonic survival in the next generation (Figure 1D). In contrast, *GFP::lem-3Y530F(241-704),* carrying the Y530F catalytic site mutation, was as hypersensitive to IR as the previously characterized *lem-3(op444)* and *lem-3(mn155)* null alleles, and the newly generated *lem-3(Δ1-704)* deletion allele which takes out the entire *lem-3* open reading frame (Figure 1D).

As previously reported for *lem-3(mn155)*, persistent bridges caused by hampering DNA replication, effected by the partial depletion of the MCM-7 subunit (via exposure to diluted RNAi), remained to a greater extent in *GFP::lem-3Y530F(241-704)* embryos compared to full length *GFP*::*lem-3(1-704)* wild-type embryos or embryos carrying *GFP::lem-3(241-704)* (Figure 1E and Figure S2A). GFP::LEM-3(241-704) and GFP::LEM-3Y530F(241-704) proteins showed the same dynamic localization as previously reported for full-length GFP::LEM-3(1-704); excluded from the nucleus in interphase, and midbody accumulation occurring during cytokinesis (Figure S2B). However, GFP::LEM-3(241-704) and GFP::LEM-3Y530F(241-704) appeared earlier, already at anaphase/ telophase, with more extensive midbody accumulation compared to full-length wild-type GFP::LEM-3(1-704) (Figure 1F, G and Figure S2B). Western blot analysis revealed that the levels of GFP::LEM-3(241-704)::3xHA and GFP::LEM-3Y530F(241-704)::3xHA were increased as compared to the levels of full-length wild-type GFP::LEM-3(1-704)::3xHA (Figure S3A, B).

Full-length GFP::LEM-3(1-704) can be detected already at anaphase/telophase by more sensitive staining of fixed embryos, indicating that there is a quantitative but no qualitative difference between GFP::LEM-3(241-704) and full-length GFP::LEM-3(1-704) localization (Figure S4, (13)) (Interestingly, we observed that a small fraction of the full-length GFP::LEM-3(1-704) protein but not the GFP::LEM-3(241-704) protein localizes to the plasma membrane (Figure S4)). We suspect that the LEM-3 N-terminal Ankyrin domain (aa26-138) restricts LEM-3 protein expression, as GFP::LEM-3(1-25,139-704) dynamic localization and protein expression levels mirrors those of GFP::LEM-3(241-704) (Figure 1F,G, S2B and S3A,B). We suspect that the Ankyrin repeat domain, directly or indirectly, contributes to LEM-3 degradation. The GFP::LEM-3(241-524, 637-704) derivative lacking the catalytic domain and conferring a null phenotype showed delayed and reduced localization to the midbody, confirming that the GIY-YIG domain is strictly required for LEM-3 function but also contributes to LEM-3 localization (Figure S2B).

All in all, these data indicate (i) that the N-terminal 240 amino acids of LEM-3 are not essential for its function *in vivo*, (ii) that the conserved Y530 residue is essential for LEM-3 function *in vitro* and *in vivo*, (iii) that LEM-3(241-704) shows nuclease activity on a synthetic HJ substrate (for detail see below), and (iv) that the ankyrin repeat prevents precocious recruitment of LEM-3 and/or restricts LEM-3 protein expression.

### LEM-3(241-704) structural conservation and catalysis

To investigate reaction mechanisms we performed sequence alignments of LEM-3 and ANKLE1 with several GIY-YIG nucleases whose crystal structures have been reported, including SLX1 (29), T4 endoII (30), UvrC (28), and Hpy188I (24) (Figure S5A). Conservation was further confirmed by AlphaFold-based high-confidence structural predictions of LEM-3 and ANKLE1 (Figure 2A,B). LEM-3 and ANKLE1 predicted structures contain the common core that is present in other GIY-YIG nucleases, consisting of the three central β-sheet strands (β1, β2, and β3) and two alpha helices (α1 and α2) (Figure 2A,B). Notably, the predicted structure of the LEM-3 GIY-YIG domain contains four additional helices, two located C-terminal to β-sheet strand β1, and the other two inserted in the core fold after helix α1 (Figure 2A). These additional structural features have been commonly observed in GIY-YIG nucleases (except for SLX1) and are believed to facilitate the binding to DNA substrates and to stabilize the overall architecture (29). SLX1 is a “minimal” GIY-YIG nuclease with a core structure markedly distinct from other family members (29). Sokolowska et al. determined the crystal structures of Hpy188I/DNA complexes and this was compared to other prokaryotic GIY-YIG nucleases (24, 29, 31). The authors proposed that the catalytic mechanism is highly conserved among GIY-YIG nuclease family members. Hydrolysis of phosphodiester bonds occurs through a nucleophilic attack of a water molecule on the scissile phosphate. Within the GIY-YIG active site, a single metal ion is believed to destabilize the substrate and remains anchored during catalysis. The proposed mechanism of phosphodiester bond hydrolysis in Hpy188I involves six amino acids (Y63, H76, R84, Y88, K73, and E149), which are highly conserved and play a critical role in the catalytic activity in GIY-YIG family members (24, 29, 31).

**Figure 2:**
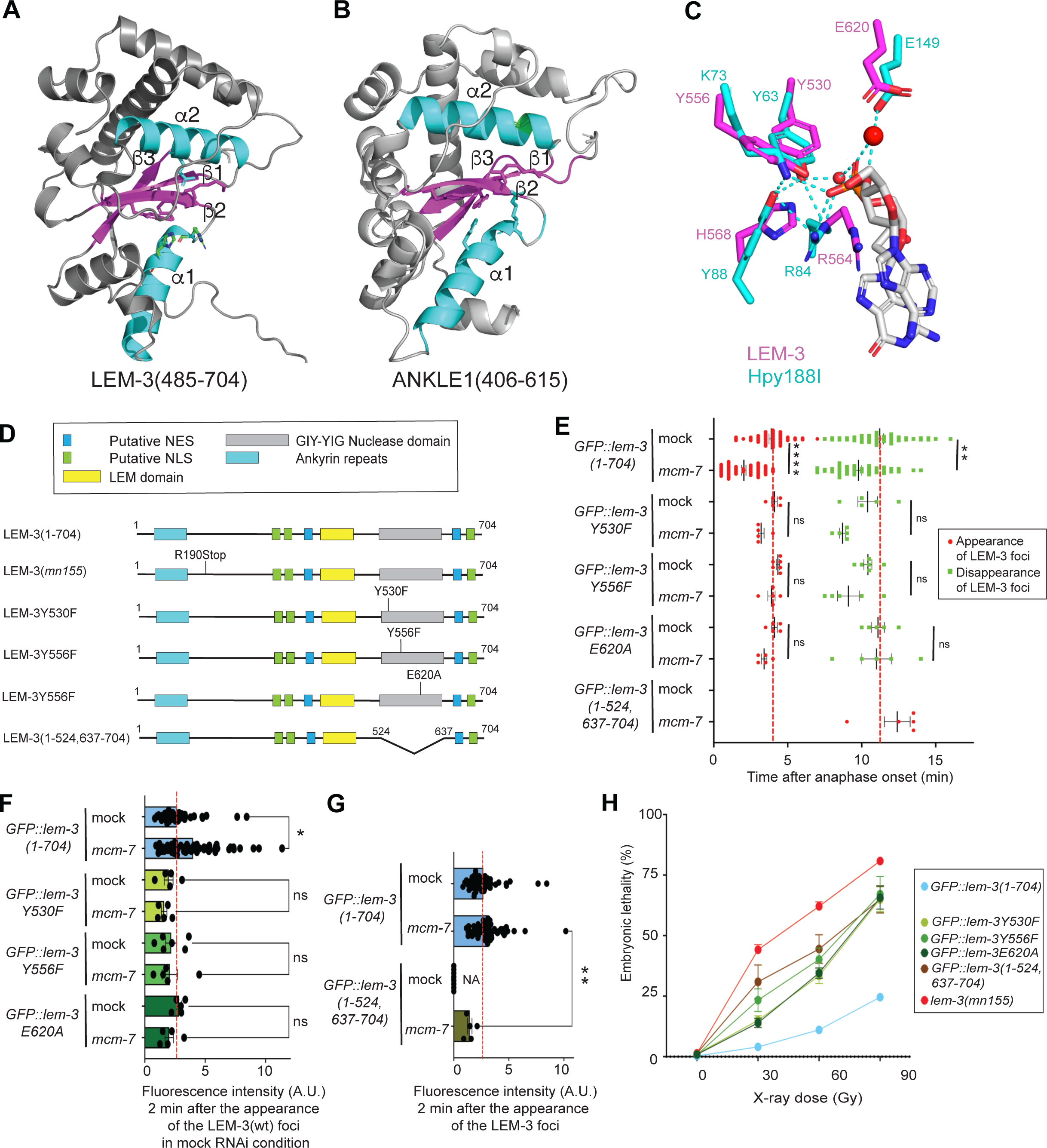
LEM-3(241-704) structural conservation and catalysis. **(A-B)** Alphafold-based high-confidence structural prediction of LEM-3(485-704) and ANKLE1 (406-615) using AlphaFold. (**C)** Overlay of active sites of LEM-3 and Hpy188I restrictase. **(D)** Schematic of the different LEM-3 derivates used in this figure. **(E)** Measurement of the time of appearance and disappearance of the LEM-3 foci for the indicated LEM-3 derivates (n≽5, mean and SEM. ns = not significant, ** p < 0.01 and **** p<0.0001 by Kruskal-Wallis with Dunn’s multiple comparison test). **(F)** The fluorescence intensity of the indicated LEM-3 derivates foci was measured 2 minutes after the appearance of the wild-type LEM-3 foci (n≽5, mean and SEM. ns = not significant and * p < 0.05 by Kruskal-Wallis with Dunn’s multiple comparison test). **(G)** The fluorescence intensity of the indicated LEM-3 derivates foci was measured 2 minutes after their appearance (n≽5, mean and SEM. ns = not significant and ** p < 0.001 by Kruskal-Wallis with Dunn’s multiple comparison test). **(H)** Analysis of the IR sensitivity of the indicated LEM-3 derivates (n≽5, mean and SEM).

To investigate the conservation of the LEM-3 catalytic mechanism, we superimposed the structure of the Hpy188I-DNA pre-cleavage complex onto the AlphaFold model of the LEM-3 GIY-YIG domain. In line with the sequence alignment, the resulting superposition of the known and predicted structures revealed a good fit, indicating a high degree of structural conservation between the known Hpy188I-DNA pre-cleavage complex and LEM-3 and ANKLE1 (Figure 2C). (i) LEM-3/ANKLE1, Y530/Y453, Y556/Y486, K559/K489, R564/R494 and H568/H498 correspond to Y63, K73, H76, R84 and Y88 in Hpy188I respectively, and are expected to participate in the activation of the water molecule. (ii) R564/R494 (R84 in Hpy188I) appears to be interacting with the 5’ oxygen atom of the scissile bond phosphate. (iii) E620/E546 (E149 in Hpy188I) is expected to coordinate the metal ion, which also interacts with the phosphate (Figure 2C). Our model agrees on hydrogen bonding between LEM-3/ANKLE1 Y530/<u>Y453</u> H568/<u>H498,</u> both residues coordinating the activating water molecule, and the proximity of Y530/<u>Y453</u> Y556/<u>Y486</u> indicating a role in acid-based catalysis (19). However, ANKLE1 N565, reported to position a metal ion to facilitate hydrolysis of the phosphate group, as well as ANKLE1 K519, are only conserved in the LEM-3/ANKLE1 eukaryotic GIY-YIG subfamily and have no core catalytic role in our model. Indeed, ANKLE1 K519 and N565 reside in an additional” helix and loop regions instead of the highly conserved core structure (Figure S5B). In summary, our modeling data suggest that the catalytic mechanism of LEM-3/ANKLE1 mirrors the mechanism observed in bacterial GIY-YIG nucleases. Residues involved in coordinating the phosphate group, the nucleophilic water molecule, and the metal ion needed for leaving group activation appear to be functionally and structurally conserved in this nuclease family.

To characterize the *in vivo* impact of catalytic site mutation in full-length *GFP::lem-3(1-704),* we generated Y530F, Y556F, and E620A mutations (Figure 2D). Substitution of catalytic residues Y530 and Y556 to F provides “atomic mutants” of LEM-3, tyrosine (Y), and phenylalanine (F) differing by only one hydroxyl group, absent in phenylalanine. We found that all catalytic mutants localized normally without *mcm-7* RNAi treatment (Figure 2E,F). As previously shown (13), wildtype LEM-3 protein localizes earlier and hyper-accumulates at the midbody upon *mcm-7* RNAi. In contrast, the midbody localization of all catalytic mutants is not advanced and increased by *mcm-7* RNAi (Figure 2E,F and Figure S6 and S7). Western-blot analysis revealed that these effects are not due to reduced levels of GFP::LEM-3Y530F(1-704)::3xHA as compared to wild-type GFP::LEM-3(1-704)::3xHA (Figure S3C). Catalytic site mutants are largely compromised; however less sensitive than the null alleles ((Figure 2H). Thus, the *in vivo* catalytic activity may not be abolished entirely and/or LEM-3 may have an additional role independent of catalysis. We generated *GFP::lem-3(1-524,637-704),* a derivate without the GIY-YIG catalytic domain, to distinguish between these possibilities. Despite the levels of GFP::LEM-3(1-524,637-704)::3xHA being slightly reduced as compared to wildtype GFP::LEM-3(1-704)::3xHA (Figure S3C), this allele also does not confer IR-hypersensitivity to the same level as *lem-3* null alleles, consistent with an auxiliary, non-catalytic role of LEM-3 (Figure 2H). The relevant function may reside in the N-terminal 240 residues of LEM-3, given that the activity of the *GFP::lem-3Y530F(241-704)* catalytic site mutant mirrors the IR-hypersensitivity of *lem-3* null alleles (Figure 1D).

Y530F, Y556F, and E620A substitutions do not cause a structural clash based on analysis in PyMol; the major predicted rotamers of the side chains of those F and A residues not interfering with the predicted structure (Figure S5C,D), and the dynamic localization of the corresponding proteins corresponding to wild type at least in the absence of *mcm-7* RNAi (Figure 2E,F). In contrast, GFP::LEM-3Y556F-G558A(1-704) (analyzed previously (13)), and GFP::LEM-3L659F(1-704), which corresponds to the *lem-3(op444)* null allele, failed to accumulate at the midbody in the first zygotic division, while low levels were observable in subsequent divisions. G558A and L659F, the latter being outwith the GIY-YIG domain, are predicted to cause steric clashes (Figure S5E,F). Western-blot analysis also revealed that the levels of GFP::LEM-3L659F(1-704)::3xHA is reduced as compared to wild-type GFP::LEM-3(1-704)::3xHA (Figure S3D), hinting that this mutant protein might have a reduced stability. Consistently, the UvrC residue G31 corresponds to LEM-3 G558, which resides in beta-sheet 2, and is positioned just behind the bound metal. Any side chains would lead to steric interference (28). In line, the corresponding G19A change in Endonuclease I-TevI curtails bacterial expression (32). All in all, catalytic site mutations block LEM-3 activity *in vivo* without overtly changing the structure of the GIY-YIG domain and affecting LEM-3 localization. LEM-3 nuclease activity is required for its role *in vivo*, the N-terminal 241 residues having a minor, non-catalytic role. In addition, the catalytic domain plays a role in LEM-3 midbody localization, catalysis likely being required for this.

### Specificity of LEM-3

We explored the substrate specificity of recombinant LEM-3(241-704) by testing its activity on various types of branched and linear DNA substrates resembling DNA repair and replication intermediates. These substrates included immobile HJ(HJ-X0-1), mobile HJ (HJ-X26), nicked HJ, replication fork (RF), 5’-flap, 3’-flap, double-stranded DNA, and single-stranded DNA, with one strand of each substrate labeled by Cy5 at the 5’ end (33, 34). Our results revealed that LEM-3(241-704) cleaves a broad range of DNA substrates (Figure 3A), and the absence of cleavage by LEM-3Y530F(241-704) serves as a control (Figure 3B). LEM-3(241-704) displayed the highest activity against 5’-flaps, 3’-flaps, and nicked HJs (details below), activity towards RFs and HJs, being ∼2-fold reduced. In contrast to the immobile HJ-X0 junction substrate, the mobile HJ (HJ-X26) carries a 26bp core at the center, allowing the point of strand exchange to migrate through those 26bp. Mobile HJs are prone to transient thermal denaturation or base pair breathing, leading to transient “bubble” structures in the junction center (35). Cleavage of the mobile HJ substrate is reduced slightly compared to the immobile HJ substrate (Figure 3A). LEM-3(241-704) activity on single-strand and double-stranded DNA substrates was weak; a smear migrating below the double-stranded DNA substrate using the highest enzyme concentrations suggests that LEM-3 may possess an additional exonuclease activity on these substrates.

**Figure 3:**
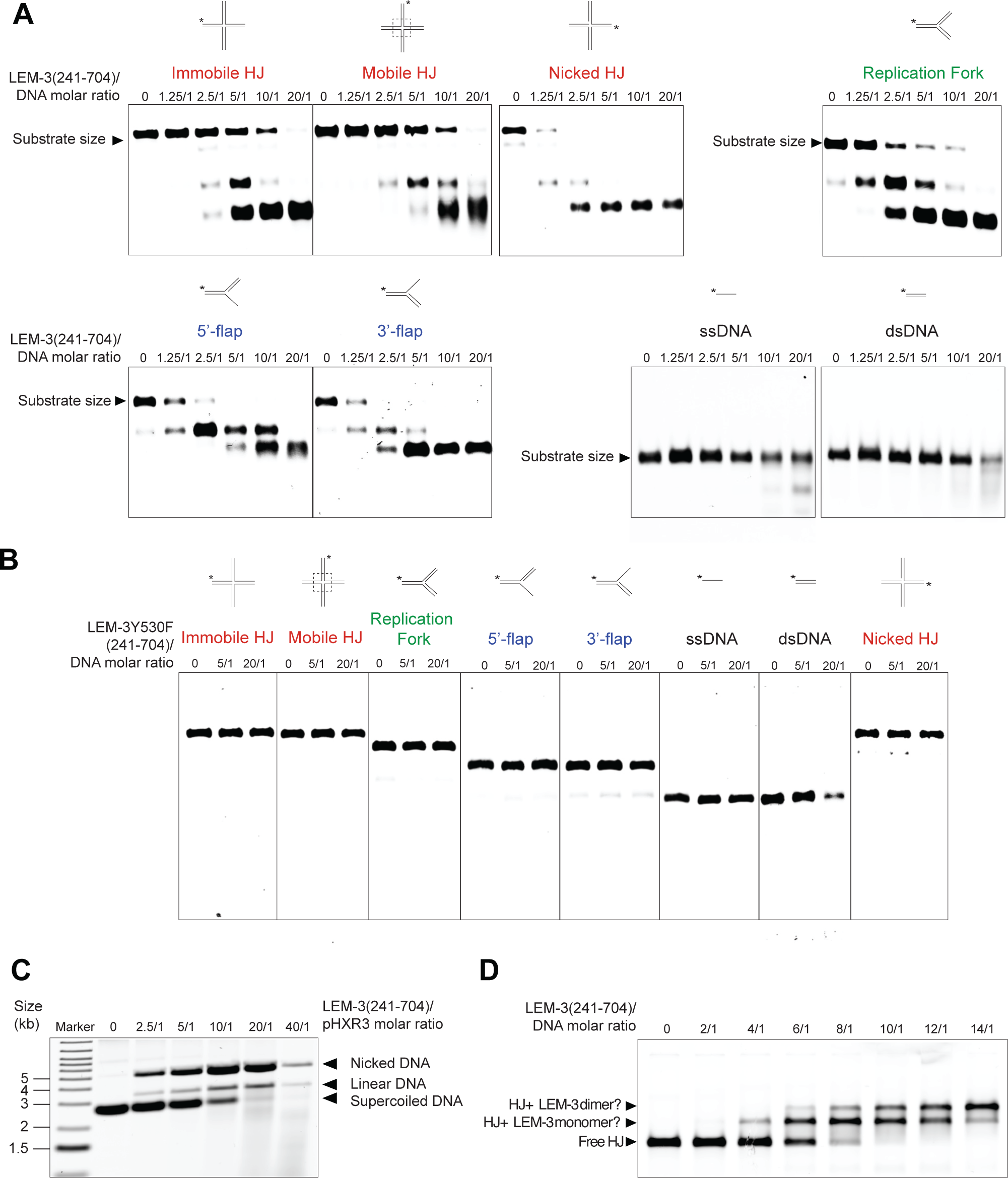
Specificity of LEM-3(241-704) and evidence of LEM-3(241-704) acting as a dimer. Nuclease activity of **(A)** LEM-3(241-704) or **(B)** LEM-3Y530F(241-704) on various model DNA substrates. Different protein/DNA substrate molar ratios were used, and incubation was performed at 37℃ for 10 mins. The cleavage products were analyzed by 6% neutral PAGE. The size of the DNA substrates is indicated. **(C)** Cruciform assay with LEM-3 (241-704). The supercoiled (uncut), the nicked (single cut), and linear (double cut) DNA were resolved by agarose gel electrophoresis. **(D)** Gel shift assay to measure the binding of LEM-3(241-704). LEM-3(241-704) was incubated for 1 hour at 4C with HJ using different protein/HJ molar ratios in the presence of 100 mM Na^+^ to prevent cleavage of HJ. The reaction was analyzed by 6% neutral PAGE. The size of the free HJ, as well as the potential HJ + LEM-3 monomer and HJ + LEM-3 dimer are indicated.

### Cleavage of HJ substrates and evidence for LEM-3 acting as a dimer

We next studied the cleavage of HJ in greater detail. While the primary sequences of the 4 branches of HJs are perfectly symmetric, junctions fold into an H-shape with two continuous strands and two crossover strands. Labeling each of the 4 strands of the HJ, we found that the HJ-X0-2 and HJ-X0-4 are cleaved slightly more efficiently, indicating that cleavage of CO strands is preferred (Figure S8A).

Cleavage of immobile and mobile junctions requires ∼5-10 higher enzyme concentration as compared to the cleavage of nicked HJ (an HJ-X0 derivative). (Figure 3A). Increased activity towards the nicked HJ substrate may indicate that cleavage on the opposite strand is enhanced upon completion of the first cleavage that generates a nicked substrate. HJ resolvases are known to bind four-way DNA junctions as homodimeric complexes, which introduce two coordinated incisions across the helical junction within the lifetime of the protein-DNA complex (36). We further examined HJ cleavage using a supercoiled pHXR3 plasmid as a substrate. This plasmid contains an inverted repeat sequence, which, upon superhelical tension, is extruded as a cruciform structure. Unilateral cleavage generates a nicked, circular DNA, whereas coordinated bilateral cleavage within the lifetime of the protein-DNA complex would produce a linear DNA product. Our results show that LEM-3(241-704) could convert pHXR3 into linear and nicked DNA at protein/DNA molar ratios from 2.5/1 to 40/1, suggesting that LEM-3(241-704) is capable of catalyzing a dual incision of the cruciform structure during the lifetime of the protein-DNA complex (Figure 3C). However, cleavage to a linear DNA substrate appears to occur at a lower rate than the formation of the nicked structure. The appearance of a smear at an enzyme to substrate concentrations higher than 10/1 indicates that, as observed for synthetic double-stranded DNA (Figure 3A), linear double-stranded DNA might be subjected to LEM-3(241-704)-dependent exonucleolytic cleavage (Figure 3C), consistent with a previous report on ANKLE1. The disappearance of the linear DNA hampers assessing if the second cleavage is accelerated in the plasmid-based cleavage assay.

To further probe the interaction between LEM-3(241-704) and HJs, we incubated LEM-3(241-704), which is rendered catalytically inactive in the presence of Na^+^, with HJ X0-1 (immobile HJ) at 4°C for 1 hour in the presence of 100mM Na^+^ and conducted mobility shift assays using 6% native/neutral PAGE. The results showed that free HJ DNA migrated as the fastest band, while protein-DNA complexes moved more slowly as two distinct protein-DNA species (Figure 3D). We postulate that the slower species likely represents a complex with a LEM-3(241-704) monomer, while the slowest species might be a dimeric LEM-3 complex bound to HJ-X0-1. This finding suggests that the protein-DNA complex might primarily form a dimer of LEM-3 bound to HJ at higher protein concentrations. Next, we determined the (K_D(DNA)_) constant of LEM-3(241-704) and HJ-X0-1. LEM-3 protein at various concentrations was incubated with 0.13nM HJ-X0-1, and the resulting products were analyzed on a native/neutral PAGE. We did not observe retarded HJ/LEM-3 complexes at lower protein concentrations (0.065nM to 17nM). In contrast, at higher protein concentrations (34nM, 67nM, and 134nM), the formation of suspected monomeric and dimeric LEM-3(241-704)/HJ complexes was detected (Figure S8B). We estimated the percentage of HJ DNA complexed by quantifying complexed and uncomplexed species and plotted as a function of LEM-3(241-704) concentration. Our results revealed that the K_D(DNA)_ of LEM-3(241-704) is ∼40nM for a DNA four-way junction, which falls within the expected range (nM-uM) of protein-DNA interactions and suggesting that LEM-3(241-704) binds to HJ with high affinity. The quantification of LEM-3(241-704) binding indicates that binding follows a hill slope, consistent with cooperative LEM-3 binding (Figure S8C). We next wished to determine if the LEM-3(241-704)/HJ-X0-1 interaction is reversible, showing that this is the case: The ratio of shifted protein observed was gradually reduced by the addition of increasing amounts of unlabeled HJ substrate (Figure S8D). All in all, we conclude that LEM-3 may form a homodimeric complex on the HJ DNA to mediate the coordinated bilateral cleavage of the junction, like other HJ-resolving enzymes such as RuvC (37, 38) and Gen1 (39– 41).

### Role of LEM-domain for catalytic activity and toxic LEM-3 derivates

To investigate the potential role of the LEM domain in LEM-3 catalytic activity, we expressed and purified a GIY-YIG domain-only protein LEM-3(485-704) (Figure S1C) and compared its activity to the activity of LEM-3(241-704). The LEM-3 ‘GIY-YIG-only’ derivate was almost entirely <u>inactive</u> on all *in vitro* DNA substrates, with residual activity only observable at the highest protein concentration (Figure 4A). The residual activity of LEM-3(485-704) and LEM-3Y530F(241-704) was higher, when the HJ extruding plasmid substrate was used, but much lower than LEM-3(241-704) (Figure 4B). These findings align with a previous publication, which showed that ANKLE1 (the human homolog of LEM-3) requires both LEM and GIY-YIG function domains for DNA cleavage *in vivo (18)*. In conclusion, our results indicate that the regions N-terminal to the GIY-YIG domain (including the LEM domain) play a critical role in efficient DNA cleavage along with the nuclease domain.

**Figure 4:**
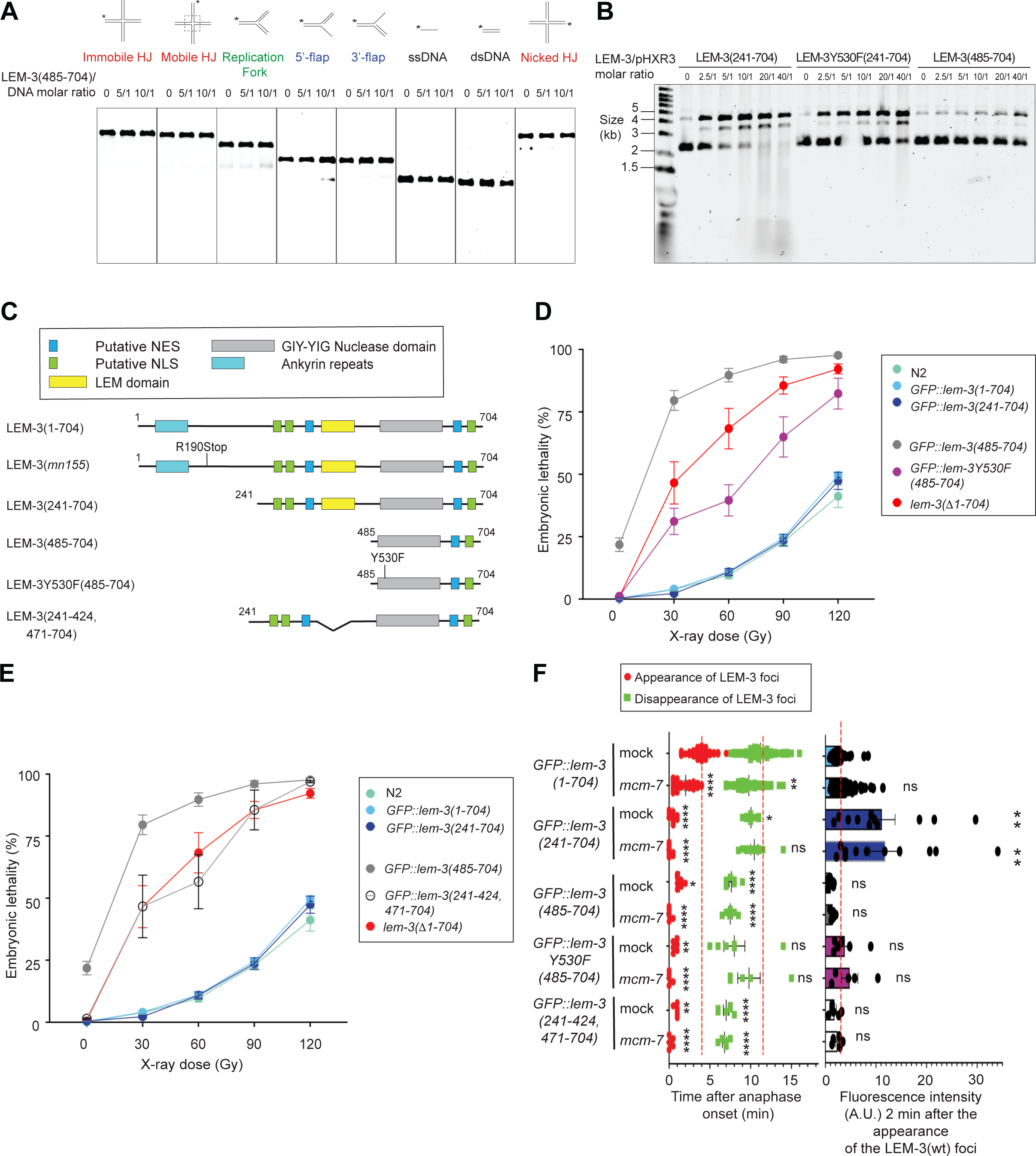
*in vitro* and *in vivo* characterization of the nuclease-only derivate LEM-3(485-704). **(A)** Nuclease activity of LEM-3(485-704) on various model DNA substrates. Different protein/DNA substrate molar ratios were used, and incubation was performed at 37℃ for 10 mins. The cleavage products were analyzed by 6% neutral PAGE. The size of the DNA substrates is indicated. **(B)** Cruciform assay with LEM-3(241-704), LEM-3Y530F(241-704) and LEM-3(485-704). The supercoiled (uncut), the nicked (single cut), and linear (double cut) DNA were resolved by agarose gel electrophoresis. **(C)** Schematic of the different LEM-3 derivates used in this figure. **(D-E)** Analysis of the IR sensitivity of the indicated LEM-3 derivates (n≽5, mean and SEM). **(F)** (Left panel) Measurement of the time of appearance and disappearance of the LEM-3 foci for the indicated LEM-3 derivates (n≽5, mean and SEM. For the time of appearance, * p < 0.05, ** p < 0.01, and **** p<0.0001 by Kruskal-Wallis with Dunn’s multiple comparison test to *GFP::lem-3(wt) mock RNAi.* For the time of disappearance, ns = not significant, * p < 0.05, ** p < 0.01 and **** p<0.0001 by Brown-Forsythe and Welch ANOVA tests with Dunnett’s T3 multiple comparisons to *GFP::lem-3(wt) mock RNAi*). (Right panel) The fluorescence intensity of the indicated LEM-3 derivates foci was measured 2 minutes after the appearance of the wild-type LEM-3 foci (n≽5, mean and SEM. ns = not significant and ** p < 0.001 by Kruskal-Wallis with Dunn’s multiple comparisons to *GFP::lem-3(wt) mock RNAi*).

Surprisingly, the strain carrying *GFP::lem-3(485-704) the* ‘GIY-YIG-only’ derivate is more sensitive to IR than *lem-3* null alleles (Figure 4C,D). In addition, even in the absence of IR treatment, progeny viability was reduced, and surviving progeny tended to grow slower than WT (Figure 4C,D). We postulate that this might be due to a failure to prevent *GFP::lem-3(485-704)* nuclear access, as evidenced by the absence of GFP::LEM-3(485-704) nuclear exclusion (evident as a halo in the wild type) observed by spinning disc microscopy (Figure S9). Consistent with such toxicity being due to nuclease activity, a strain carrying the *GFP::lem-3Y530F(485-704)* allele behaved like a null allele upon IR and did not show reduced progeny survival in the absence of IR. Thus, when nuclear, even a strongly reduced catalytic activity may be sufficient for degrading DNA. Indeed, forced nuclear expression of ANKLE1 was shown to be toxic (18). All in all, preventing LEM-3 nuclear access appears important to prevent cleavage of branched DNAs that occur during normal DNA metabolism. We note that LEM-3 contains several predicted sequences that may direct nuclear export and import (Figure 4C), of which one export site (aa 365-379) was absent in the *GFP::lem-3(485-704)* GIY-YIG only derivate.

Interestingly, *in vivo* the derivate with only the LEM domain deleted, GFP::LEM-3(241-424,471-704), was as defective as (but not more defective than) the null allele (Figure 4E). GFP::LEM-3(485-704) which only contains the GIY-YIG domain and GFP::LEM-3(241-424, 471-704) which lacks the LEM domain appears early at the spindle midzone, even before it congresses to the midbody. The level at the midbody was reduced compared to GFP::LEM-3(241-704), while the overall protein level as assessed by Western blotting was similar to GFP::LEM-3(241-704) (Figure 4F and S3E).

### Role of LEM-domain in DNA binding

We next wished to test whether the LEM-3 LEM domain has a role in DNA binding. LEM domains (LAP2-Emerin-MAN1) carry a common helix-loop-helix motif (42, 43). The canonical LEM domain mediates interaction with chromatin through binding to Barrier-to-Autointegration factor (BAF), an essential chromatin-associated protein, through hydrophobic residues on the surface of the LEM domain (44, 45). (46). Interestingly, lamina-associated polypeptide 2 (LAP2) contains two structurally divergent LEM domains, LEM and LEM-like. The LEM-like domain of LAP2 is believed to have changed during evolution in its surface residues to adapt for DNA binding (45). LEM, LEM-like, and SAP domains are evolutionarily related motifs that exhibit a remarkably similar structural organization with an N-terminal three-residue helical turn and two helices connected by an extended loop (47). The SAP domain of SLX4, a scaffold protein that coordinates the action of several nucleases including SLX1, can also bind to DNA through its positively charged residues (29). We, therefore, hypothesize that the LEM domain of LEM-3 is directly involved in DNA binding through positively charged surface residues, as observed in the LEM-like motif of LAP2 and the SLX-4 SAP domain (29),

We utilized AlphaFold2 to predict the structure of LEM-3(241-704), and PyMOL to analyze the electrostatic potential distribution (Figure 5A,B). The predicted LEM-3(241-704) LEM domain shares a conserved structure with unique features that differ from other LEM motifs, and we consider it (and from now on refer to it) as LEM-like for the following reasons. It is predicted to comprise three helices, a short helix in the N-terminus, two long helices corresponding to the helical turn, and helix1 and 2 found in LEM and LEM-like domains of LAP2 (Figure 5A). Importantly, we identified a positively charged surface in the LEM-3 LEM domain rather than a hydrophobic surface (Figure 5B). Indeed, comparing the LEM-3 AlphaFold model to the crystal structure of the *S. cerevisiae* SLX1/SLX4 complex bound to a 5’flap (the SLX4 fragment resolved, containing the SAP and a CCD domain, required for SLX1 binding (29)) reveals that LEM-3(241-704) and SLX1–SLX4(SAP+CCD) share structural similarities (Figure 5A). Both contain a conserved GIY-YIG catalytic domain, as well as two open, positively charged surface areas, one in the GIY-YIG domain and one in the LEM-like (LEM-3) and the SAP domains (SLX4), respectively (23, 24) (Figure 5B) (47). Consistent with a role of the LEM-3 LEM-like domain in DNA binding, structure prediction using AlphaFold3 confirmed the aforementioned bipartite DNA binding mode of LEM-3, both the LEM-like and GIY-YIG domains binding to DNA (Figure 5H).

**Figure 5:**
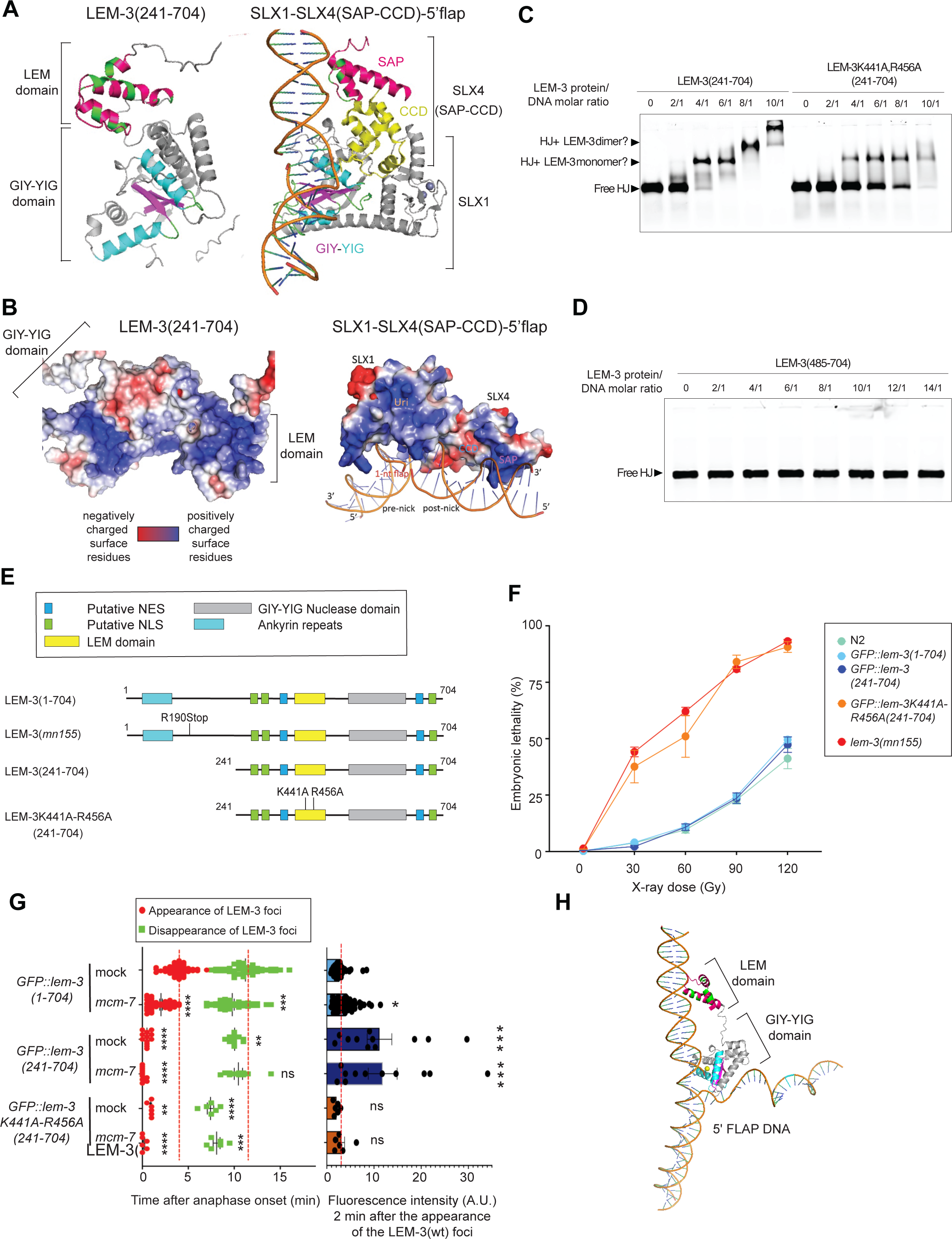
The LEM domain of LEM-3 contributes to its DNA binding *in vitro* and its midbody localization and activity *in vivo*. **(A)** AlphaFold prediction of the structure of LEM-3(241-704) compared to the crystal structure of the *S. cerevisiae* SLX1/SLX-4 complex bound to a 5’flap (29). **(B)** PyMOL analysis of the electrostatic potential distribution of LEM-3(241-704) compared to the *S. cerevisiae* SLX1/SLX-4 complex bound to a 5’-flap (29). **(C)** Gel shift assay to measure the binding of LEM-3(241-704) or LEM-3K441A;R456A(241-704) to HJ. LEM-3(241-704) or LEM-3K441A;R456A(241-704) were incubated for 1 hour at 4℃ with HJ using different protein/HJ molar ratio and in the presence of 100mM Na^+^ to prevent cleavage of HJ. The reaction was analyzed by 6% neutral PAGE. The size of the free HJ, as well as the potential HJ + LEM-3 monomer and HJ + LEM-3 dimer are indicated. **(D)** Gel shift assay to measure the binding of LEM-3(484-704) to HJ. LEM-3(484-704) was incubated for 1 hour at 4C with HJ using different protein/HJ molar ratios in the presence of 100 mM Na^+^ to prevent cleavage of HJ. The reaction was analyzed by 6% neutral PAGE. The size of the free HJ is indicated. **(E)** Schematic of the different LEM-3 derivates used in this figure. **(F)** Analysis of the IR sensitivity of the indicated LEM-3 derivates (n≽5, mean and SEM). **(G)** (Left panel) Measurement of the time of appearance and disappearance of the LEM-3 foci for the indicated LEM-3 derivates (n≽5, mean and SEM. For the time of appearance, ** p < 0.01 and **** p<0.0001 by Kruskal-Wallis with Dunn’s multiple comparison test to *GFP::lem-3(wt) mock RNAi.* For the time of disappearance, ns = not significant, ** p < 0.01, *** p < 0.001 and **** p<0.0001 by Brown-Forsythe and Welch ANOVA tests with Dunnett’s T3 multiple comparisons to *GFP::lem-3(wt) mock RNAi*). (Right panel) The fluorescence intensity of the indicated LEM-3 derivates foci was measured 2 minutes after the appearance of the wild-type LEM-3 foci (n≽5, mean and SEM. ns = not significant, * p < 0.05 and *** p < 0.001 by Kruskal-Wallis with Dunn’s multiple comparisons to *GFP::lem-3(wt) mock RNAi*). **(H)** AlphaFold3 structure prediction of LEM-3 with 5’ Flap.

To find critical residues required for DNA binding, we conducted sequence alignments and analyzed the predicted structure of the LEM-3 (Figure S10). We narrowed our attention to. K441 and R456, residues located on helix 2, and the N-terminus of helix 3 in the LEM-3 LEM-like domain and aligned with the proposed DNA interaction surfaces in helix 1 and the N-terminus of helix 2 of the LAP2 domain (Figure S10). Next, we tested the ability of LEM-3K441A-R456A(241-704) to bind HJ using electrophoretic mobility shift assays (Figure S1D, Figure 5C). The LEM-3K441A-R456A(241-704) double mutant showed significantly weaker substrate binding compared to the wild-type LEM-3(241-704), with residual binding activity in contrast to LEM-3(485-704), which lacks the LEM-like domain (Figure 5D). Consistent with reduced DNA binding, the *GFP::lem-3K441A-R456A(241-704)* allele is hypersensitive to IR (Figure 5E,F). GFP::LEM-3K441A-R456A(241-704), like GFP::LEM-3(241-704), appears early at the spindle midzone, with reduced levels at the midbody compared to GFP::LEM-3(241-704). (Figure 5E,G and S3F). All in all, these data indicate that the LEM-3 LEM-like domain aids nuclease activity *in vivo* and *in vitro*, contributes to preventing LEM-3 mis-localization, and is required for strong midbody localization, and/or the prevention of premature midbody dissociation.

Nevertheless, our *in vivo* data indicate that the LEM-3 N-terminus, encompassing the Ankyrin repeat domain, carries a function redundant with the LEM-like domain, as ‘full-length’ *GFP::lem-3(1-425, 470-704)* carrying a deletion of the LEM-like domain and the corresponding ‘full-length’ *GFP::lem-3K441A-R456A(1-704)* allele show an intermediate level of IR sensitivity (as opposed *GFP::lem-3K441A-R456A(241-704),* which is as sensitive as the *lem-3* null alleles (Figure 6A,B)), even so, full-length proteins are less abundant than GFP::LEM-3(241-704) as determined by Western blotting (Figure S3G). Consistent with the redundant role of the N-terminal Ankyrin repeat domain, GFP::LEM-3(1-425, 470-704) *and* GFP::LEM-3K441A-R456A(1-704) midbody association is reduced and occurs with delayed kinetics compared to wild-type GFP::LEM-3(1-704), a pattern also observed in embryos carrying GFP::LEM-3(1-425), a derivative that only carries residues N-terminal of the LEM-like domain (Figure 6C,D). Analyzing GFP::LEM-3(1-425), which solely carries residues N-terminal to the LEM-like domain (and is as IR defective as the *lem-3* null alleles), we found that midbody association is absent without *mcm-7* RNAi treatment and occurs with delayed kinetics compared to wild-type *GFP::LEM-3(1-704*) in the presence of partial *mcm-7* RNAi (Figure 6C,D). These results confirm that the LEM-3-like domain and amino acids predicted to be involved in facilitating DNA binding within the LEM-like domain are essential for normal LEM-3 localization, also in the context of the full-length protein.

**Figure 6:**
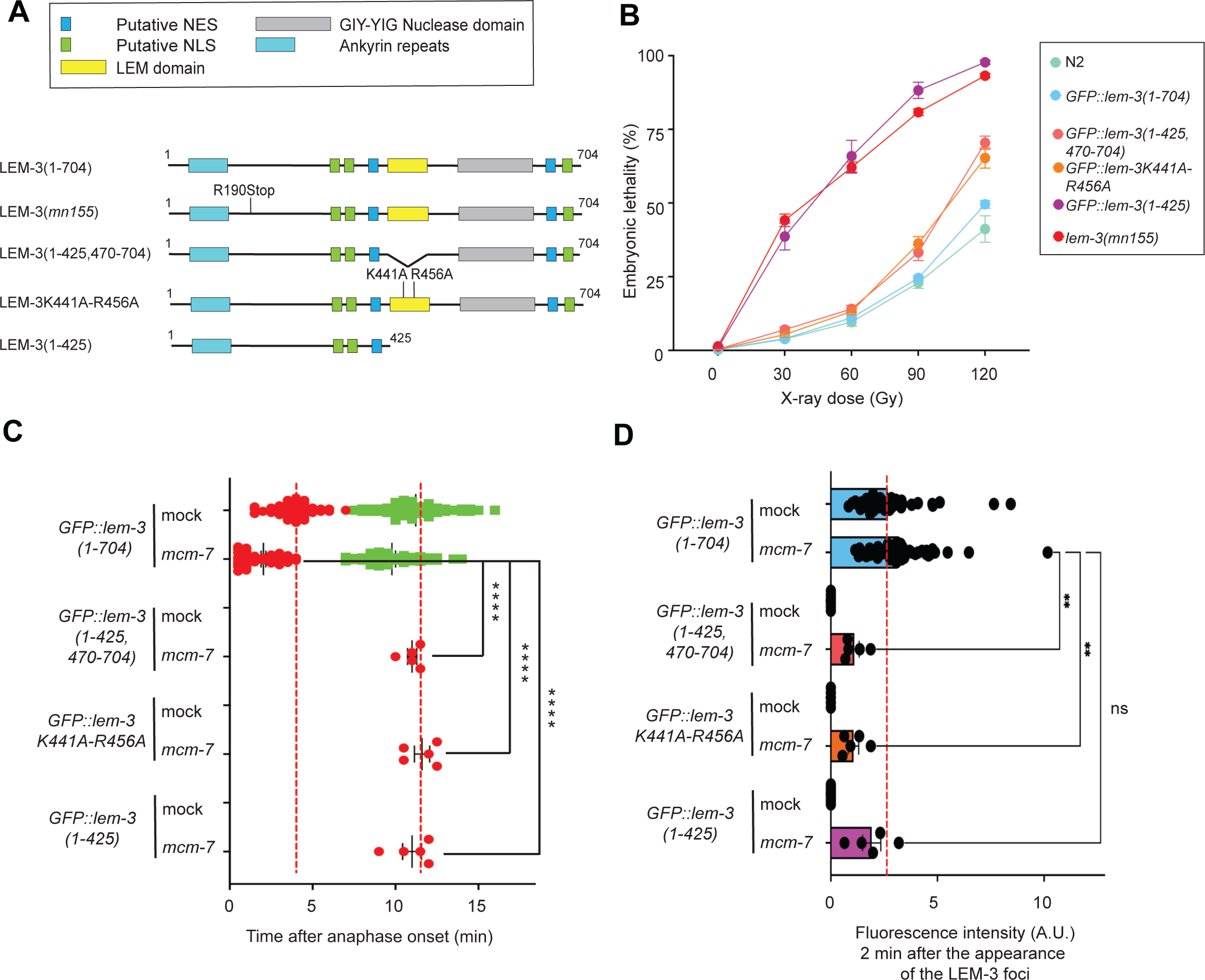
Role of the N-terminal domain of LEM-3. **(A)** Schematic of the different LEM-3 derivates used in this figure. **(B)** Analysis of the IR sensitivity of the indicated LEM-3 derivates (n≽5, mean and SEM). **(C)** Measurement of the time of appearance and disappearance of the LEM-3 foci for the indicated LEM-3 derivates (n≽5, mean and SEM. **** p<0.0001 by Kruskal-Wallis with Dunn’s multiple comparison test to *GFP::lem-3(wt) mcm-7 RNAi.*) **(D)** The fluorescence intensity of the indicated LEM-3 derivates foci was measured 2 minutes after the appearance of the wild-type LEM-3 foci (n≽5, mean and SEM. ns = not significant, ** p < 0.01 by Kruskal-Wallis with Dunn’s multiple comparisons to *GFP::lem-3(wt) mcm-7 RNAi*).

## Discussion

It is pivotal for cells to process DNA bridges just before cells divide. We found that the purified LEM-3 protein can cleave a wide range of branched DNA species *in vitro,* consistent with LEM-3 acting as a ‘last chance catch-all’ enzyme that processes DNA bridges that can originate from persistent recombination intermediates, local DNA under-replication, and chromosome entanglement (13). DNA flaps, nicked HJ and HJ result from remaining intermediates of recombinational repair. In our experimental system of *C. elegans* first zygotic cell division, bridges also arise from persistent intermediates formed during preceding meiotic recombination. Some of these diverse branched DNA structures are expected to remain, if not processed prior to cytokinesis progression by the BTR complex during the S-phase, the SMX tri-nucleases complex in G2, and the GEN1 nuclease in anaphase (48). The redundancy and step-by-step action of these HJ junction processing activities, where LEM-3/ANKLE1 provides the ‘last chance catch-all’ activity, is a testament to the immense importance of processing all DNA bridges before cells divide. Such redundancy might explain why ANKLE1 has only mild phenotypes in human cells (14), the prior mechanisms being highly effective, and cell division taking much longer in mammals compared to nematode embryonic cell divisions.

Well-characterized RuvC-like HJ-resolving enzymes catalyze HJ resolution in bacteria. These cleave HJ in perfect symmetry and act as dimers facilitating HJ cleavage by a nick, counter-nick mechanism, with the second cleavage facilitated by the first nick (37, 38). Based on this precedent, studies on GEN1 initially focused on its HJ activity (49). Nevertheless, GEN1 and its yeast ortholog Yen1, like LEM-3/ANKLE1, also cleave further substrates (50, 51). The same applies to the enzymes associated with the SMX tri-nucleases complex, where MUS81, like LEM-3, preferably cleaves, nicked HJs and flap structures (52). All in all, mammalian HJ processing enzymes appear remarkably promiscuous in their substrate selection compared to their bacterial counterparts, unrelated by primary sequence. Nevertheless, HJs are cleaved by nick counter-nick mechanics, favored by HJ substrate-mediated GEN1 dimerization, or SLX4/SLX1 nickage, followed by MUS81 counter-nickage of the nicked HJ substrate. Our data point towards LEM-3 possibly cleaving HJ based on a reaction mechanism akin to the mechanism shown for GEN1. A recent study did not find evidence for ANKLE1 dimerization on HJ substrates or coordinated cleavage (19), but complete cleavage of an extruded HJ from a supercoiled plasmid was shown (14, 18). High LEM-3 and ANKLE1 levels or long reaction times reveal an additional nucleolytic activity leading to the degradation of the substrate, thereby complicating the determination of the reaction kinetics of cleavage and counter cleavage (14, 18).

Based on our previous genetic data, LEM-3 also has a role in helping to safeguard unreplicated DNA through cytokinesis, a feat only possible when the single double-stranded DNA is separated into two single strands. To achieve this, LEM-3 might take advantage of its nickase activity, which functionally could be akin to the first step of a topoisomerase reaction. We did not observe nickage of our double-stranded *in vitro* oligonucleotide substrate, but nickage occurs on plasmid substrates due to the extrusion of branched secondary structures. Such structures readily form on native DNA as a consequence of torsal tension resulting from topoisomerase activity.

The promiscuity of nuclease activities raises a most important and, in our opinion, largely unresolved question, which is how the very same substrates cleaved by GEN1, SMX, and LEM-3/ANKLE1 are prevented from being cleaved when they arise during replication, ongoing DNA repair, or upon torsional stress associated with supercoiling. Indeed, there is evidence that both GEN1 and XPF1 cause mutagenesis by cleaving endogenous DNA, which is predicted to form HJ or loop-like structures *in vivo* (53–55). We know very little of how this is achieved. For GEN1 it was shown that it is excluded from the nucleus in mammalian cells to restrict its function to mitosis. Intriguingly, when the catalytic domain of LEM-3 is expressed, this fragment is toxic, even though it is largely (but not fully) inactive *in vitro*. We postulate that remnant activity conferred this toxicity, as rendering the LEM-3 catalytic domain only fragment inactive suppresses this effect. LEM-3(485-704) lacks a predicted NES sequence N-terminal of the LEM domain (aa 365-379). Interestingly, even though these NES are not conserved at the sequence level, the NES that mediates ANKLE1 nuclear export is also N-terminal of the LEM domain, and in the absence of this NES nuclear ANKLE1 confers genotoxicity (56). LEM-3(485-704) contains one predicted NLS at the C-terminus of the protein.

We found that the LEM domain of LEM-3 has at least two functions. Canonical LEM domains mediate association with the nuclear envelope protein BAF via hydrophobic interactions (46). Interestingly, LEM-3(485-704), which lacks the LEM domain, does not bind HJ and is largely inactive on any of the branched DNA substrates. Since the LEM-3 LEM domain contains conserved positively charged residues on its surface, we think that the LEM domain of LEM-3 is a LEM-like domain that contributes to the binding of LEM-3 to its DNA substrate, a hypothesis in line with the DNA binding capacity of LAP2. Indeed, our data indicate that the bipartite structure of LEM-3, comprised of a LEM-like DNA binding domain and a GIY-YIG catalytic domain functions akin to SLX-1 composed mainly of the GIY-YIG in conjunction with SLX-4 where the embedded LEM domain confers DNA binding.

The LEM-3 LEM-like domain also contributes to LEM-3 midbody localization. This contrasts with recent observations in mammalian cells, where ANKLE1’s LEM domain is dispensable for midbody localization. Instead, ANKLE1 midbody localization requires its Ankyrin Repeat domain (14). How the LEM-3 LEM-like domain contributes to its midbody localization remains to be elucidated. The LEM-like domain might mediate LEM-3 interaction with midbody proteins. Alternatively, LEM-3 might bind DNA through its LEM-like domain, and this binding to DNA might be necessary to target LEM-3 in the midbody. Consistent with this latter scenario, the LEM-3 GIY-YIG nuclease domain, which is also essential for DNA binding, plays a vital role in LEM-3 midbody localization. The nematode LEM-3 Ankyrin domain and or residues immediately downstream thereof are required to restrict protein levels, leading to a ∼3-fold increase in LEM-3. We postulate that this is likely due to ubiquitin-mediated protein degradation. Interestingly, this LEM-3(241-704) derivate appears at the spindle midzone already before LEM-3 congresses to the midbody, consistent with our previous observations of full-length LEM-3 when observed by most sensitive immunolocalization showing the same pattern. In line with this, LEM-3 might indeed already start to act in anaphase, concomitant with GEN1, and by inference, LEM-3 midbody localization might be the final stage of LEM-3 function. Intriguingly, we only observed LEM-3 phosphorylation at the midbody and not before midbody congression (13), although experiments have to be taken with caution as the sensitivity of the LEM-3 phospho-antibody may be too low for detecting more dispersed LEM-3 protein.

Indeed, the regulation of LEM-3 localization is more intricate. We previously showed that the AIR-2 Aurora kinase phosphorylates LEM-3 at residues (S194 and S196) embedded in the unstructured area between the N-terminal Ankyrin repeat domain and the central LEM-like domain. Mutating these phosphorylation sites largely attenuates LEM-3 midbody localization and function. We considered these sites being conserved with ANKLE1, but this is not the case; as a previously predicted exon is no longer included in the actual ANKLE1 cDNA. Intriguingly, when we deleted the entire unstructured region more, we observed that LEM-3 functions normally. It will be interesting to work out details as to how this phosphorylation functions. Unstructured domains often acquire a structure upon binding to an interacting protein, and we postulate this might also be the case for LEM-3. ANKLE1 does not carry this phosphorylation site, but ANKLE1 localization does not depend on the Aurora B kinase.

Our results suggest that LEM-3 also has a non-catalytic function, as mutating conserved catalytic residues or removing the entire catalytic domain largely but not fully mimics the hypersensitivity of the *lem-3* null allele. At present, we can only speculate about these functions. We failed to observe a difference in the timing of cytokinesis in the absence of *lem-3,* cell cycles being extremely fast during the first mitotic divisions, and the exact point of cytokinesis completion being hard to discern. It will be interesting to test if the timing of later postembryonic divisions is propagated without *lem-3* and if this also involves non-catalytic functions. Intriguingly, we observed that full-length LEM-3 is associated with the plasma membrane. While the functional relevance of this subcellular localization remains elusive, it requires the N-terminus of the LEM-3 protein, as GFP::LEM-3(241-704) does not localize to the plasma membrane.

All in all, our data provide important insights as to how LEM-3 acts to safeguard genome integrity and prevents aneuploidy by processing DNA bridges just before cells divide. We show that basic core reaction mechanisms are conserved from bacteria to animals, but details of how LEM-3 and ANKLE1 are regulated will likely differ. Nevertheless, both enzymes, based on the increased sensitivity of knockouts, much more so in the worm, contribute to genome stability. It remains to be investigated how this tallies with the inherent toxicity of cleaving late DNA bridges, which is predicted to cause toxic chromosome-to-chromosome fusions, and ensuring cellular dysfunction in the following divisions. It will be interesting to work out further how LEM-3 and ANKLE1 preserve genome integrity.

## Acknowledgments

We want to thank the members of the Gartner laboratory and the Korean Institute for Basic Science Center for Genomic Integrity for their fruitful discussions. We are also grateful to members of the Dundee School of Life Sciences, particularly to Axel Knebel, Elisa Garcia-Wilson, Nicola Wiechens, James Hastie for protein purification and advice with native gels. We also thank Orlando Schaerer and Nadin Memar for their comments on the manuscript. We thank Luthfiyyah Mutsnaini for her excellent technical support. We thank Prof KJ Myung for his unwavering support.

## Author Contributions

J.S. performed the *in vitro* experiments (LEM-3 protein purification, nuclease assays, EMSA, and AlphaFold prediction) and contributed to the writing of the *in vitro* part of the manuscript. P.G. performed the *C.elegans in vivo* experiments (IR sensitivity assay, partial *lem-3* sensitivity assay, spinning disk imaging, and immunostaining experiments) and contributed to the writing of the *in vivo* part of the manuscript. S.G.M.R generated some of the transgenic animals as indicated in Table S1 and performed Western-Blot analyses.

Y.H. generated strains and contributed to the writing of the manuscript, J. S, S.G.M.R and A.G. designed the experiments and wrote the manuscript. A.G secured funding (BB/S002782/1).

## References

1. Finardi, A., Massari, L.F. and Visintin, R. (2020) Anaphase Bridges: Not All Natural Fibers Are Healthy. Genes, 11.

2. McClintock, B. (1941) The Stability of Broken Ends of Chromosomes in Zea Mays. Genetics, 26, 234–282.

3. Liu, Y., Nielsen, C.F., Yao, Q. and Hickson, I.D. (2014) The origins and processing of ultra fine anaphase DNA bridges. Curr. Opin. Genet. Dev., 26, 1–5.

4. Shi, Q. and King, R.W. (2005) Chromosome nondisjunction yields tetraploid rather than aneuploid cells in human cell lines. Nature, 437, 1038–1042.

5. Fujiwara, T., Bandi, M., Nitta, M., Ivanova, E.V., Bronson, R.T. and Pellman, D. (2005) Cytokinesis failure generating tetraploids promotes tumorigenesis in p53-null cells. Nature, 437, 1043–1047.

6. Hoffelder, D.R., Luo, L., Burke, N.A., Watkins, S.C., Gollin, S.M. and Saunders, W.S. (2004) Resolution of anaphase bridges in cancer cells. Chromosoma, 112, 389–397.

7. Steigemann, P., Wurzenberger, C., Schmitz, M.H.A., Held, M., Guizetti, J., Maar, S. and Gerlich, D.W. (2009) Aurora B-mediated abscission checkpoint protects against tetraploidization. Cell, 136, 473–484.

8. Nazaryan-Petersen, L., Bjerregaard, V.A., Nielsen, F.C., Tommerup, N. and Tümer, Z. (2020) Chromothripsis and DNA Repair Disorders. J. Clin. Med. Res., 9.

9. Hong, Y., Zhang, H. and Gartner, A. (2021) The Last Chance Saloon. Front Cell Dev Biol, 9, 671297.

10. Guervilly, J.-H. and Gaillard, P.H. (2018) SLX4: multitasking to maintain genome stability. Crit. Rev. Biochem. Mol. Biol., 53, 475–514.

11. Matos, J., Blanco, M.G., Maslen, S., Skehel, J.M. and West, S.C. (2011) Regulatory control of the resolution of DNA recombination intermediates during meiosis and mitosis. Cell, 147, 158–172.

12. Chan, Y.W. and West, S.C. (2014) Spatial control of the GEN1 Holliday junction resolvase ensures genome stability. Nat. Commun., 5, 4844.

13. Hong, Y., Sonneville, R., Wang, B., Scheidt, V., Meier, B., Woglar, A., Demetriou, S., Labib, K., Jantsch, V. and Gartner, A. (2018) LEM-3 is a midbody-tethered DNA nuclease that resolves chromatin bridges during late mitosis. Nat. Commun., 9, 728.

14. Jiang, H., Kong, N., Liu, Z., West, S.C. and Chan, Y.W. (2023) Human Endonuclease ANKLE1 Localizes at the Midbody and Processes Chromatin Bridges to Prevent DNA Damage and cGAS-STING Activation. Adv. Sci., 10, e2204388.

15. Sonneville, R., Querenet, M., Craig, A., Gartner, A. and Blow, J.J. (2012) The dynamics of replication licensing in live Caenorhabditis elegans embryos. J. Cell Biol., 196, 233–246.

16. Edgar, L.G. and McGhee, J.D. (1988) DNA synthesis and the control of embryonic gene expression in C. elegans. Cell, 53, 589–599.

17. Braun, J., Meixner, A., Brachner, A. and Foisner, R. (2016) The GIY-YIG Type Endonuclease Ankyrin Repeat and LEM Domain-Containing Protein 1 (ANKLE1) Is Dispensable for Mouse Hematopoiesis. PLoS One, 11, e0152278.

18. Brachner, A., Braun, J., Ghodgaonkar, M., Castor, D., Zlopasa, L., Ehrlich, V., Jiricny, J., Gotzmann, J., Knasmüller, S. and Foisner, R. (2012) The endonuclease Ankle1 requires its LEM and GIY-YIG motifs for DNA cleavage in vivo. J. Cell Sci., 125, 1048–1057.

19. Freeman, A.D.J., Déclais, A.-C., Wilson, T.J. and Lilley, D.M.J. (2023) Biochemical and mechanistic analysis of the cleavage of branched DNA by human ANKLE1. Nucleic Acids Res., 51, 5743–5754.

20. Song, J., Freeman, A.D.J., Knebel, A., Gartner, A. and Lilley, D.M.J. (2020) Human ANKLE1 Is a Nuclease Specific for Branched DNA. J. Mol. Biol., 432, 5825–5834.

21. Maciejowski, J., Li, Y., Bosco, N., Campbell, P.J. and de Lange, T. (2015) Chromothripsis and Kataegis Induced by Telomere Crisis. Cell, 163, 1641–1654.

22. Przanowski, P., Przanowska, R.K. and Guertin, M.J. (2023) ANKLE1 cleaves mitochondrial DNA and contributes to cancer risk by promoting apoptosis resistance and metabolic dysregulation. Commun Biol, 6, 231.

23. Dunin-Horkawicz, S., Feder, M. and Bujnicki, J.M. (2006) Phylogenomic analysis of the GIY-YIG nuclease superfamily. BMC Genomics, 7, 1–19.

24. Sokolowska, M., Czapinska, H. and Bochtler, M. (2011) Hpy188I-DNA pre- and post-cleavage complexes--snapshots of the GIY-YIG nuclease mediated catalysis. Nucleic Acids Res., 39, 1554–1564.

25. Edgell, D.R. and Shub, D.A. (2001) Related homing endonucleases I-*Bmo*I and I-*Tev*I use different strategies to cleave homologous recognition sites. Proc. Natl. Acad. Sci. U. S. A., 98, 7898–7903.

26. Carlson, K. and Wiberg, J.S. (1983) In vivo cleavage of cytosine-containing bacteriophage T4 DNA to genetically distinct, discretely sized fragments. J. Virol., 48, 18–30.

27. Bujnicki, J.M., Radlinska, M. and Rychlewski, L. (2001) Polyphyletic evolution of type II restriction enzymes revisited: two independent sources of second-hand folds revealed. Trends Biochem. Sci., 26, 9–11.

28. Truglio, J.J., Rhau, B., Croteau, D.L., Wang, L., Skorvaga, M., Karakas, E., DellaVecchia, M.J., Wang, H., Van Houten, B. and Kisker, C. (2005) Structural insights into the first incision reaction during nucleotide excision repair. EMBO J., 24, 885–894.

29. Xu, X., Wang, M., Sun, J., Yu, Z., Li, G., Yang, N. and Xu, R.-M. (2021) Structure specific DNA recognition by the SLX1-SLX4 endonuclease complex. Nucleic Acids Res., 49, 7740–7752.

30. Andersson, C.E., Lagerbäck, P. and Carlson, K. (2010) Structure of bacteriophage T4 endonuclease II mutant E118A, a tetrameric GIY-YIG enzyme. J. Mol. Biol., 397, 1003– 1016.

31. Ibryashkina, E.M., Zakharova, M.V., Baskunov, V.B., Bogdanova, E.S., Nagornykh, M.O., Den’mukhamedov, M.M., Melnik, B.S., Kolinski, A., Gront, D., Feder, M., et al. (2007) Type II restriction endonuclease R.Eco29kI is a member of the GIY-YIG nuclease superfamily. BMC Struct. Biol., 7, 48.

32. Kowalski, J.C., Belfort, M., Stapleton, M.A., Holpert, M., Dansereau, J.T., Pietrokovski, S., Baxter, S.M. and Derbyshire, V. (1999) Configuration of the catalytic GIY-YIG domain of intron endonuclease I-TevI: coincidence of computational and molecular findings. Nucleic Acids Res., 27, 2115–2125.

33. Wyatt, H.D.M., Sarbajna, S., Matos, J. and West, S.C. (2013) Coordinated actions of SLX1-SLX4 and MUS81-EME1 for Holliday junction resolution in human cells. Mol. Cell, 52, 234– 247.

34. Wyatt, H.D.M., Laister, R.C., Martin, S.R., Arrowsmith, C.H. and West, S.C. (2017) The SMX DNA Repair Tri-nuclease. Mol. Cell, 65, 848–860.e11.

35. Ciccia, A., Constantinou, A. and West, S.C. (2003) Identification and Characterization of the Human Mus81-Eme1 Endonuclease*. J. Biol. Chem., 278, 25172–25178.

36. West, S.C., Blanco, M.G., Chan, Y.W., Matos, J., Sarbajna, S. and Wyatt, H.D.M. (2015) Resolution of Recombination Intermediates: Mechanisms and Regulation. Cold Spring Harb. Symp. Quant. Biol., 80, 103–109.

37. Dunderdale, H.J., Benson, F.E., Parsons, C.A., Sharpies, G.J., Lloyd, R.G. and West, S.C. (1991) Formation and resolution of recombination intermediates by E. coliRecA and RuvC proteins. Nature, 354, 506–510.

38. Dunderdale, H.J., Sharples, G.J., Lloyd, R.G. and West, S.C. (1994) Cloning, overexpression, purification, and characterization of the Escherichia coli RuvC Holliday junction resolvase. J. Biol. Chem., 269, 5187–5194.

39. Freeman, A.D.J., Liu, Y., Déclais, A.-C., Gartner, A. and Lilley, D.M.J. (2014) GEN1 from a thermophilic fungus is functionally closely similar to non-eukaryotic junction-resolving enzymes. J. Mol. Biol., 426, 3946–3959.

40. Liu, Y., Freeman, A.D.J., Déclais, A.-C., Wilson, T.J., Gartner, A. and Lilley, D.M.J. (2015) Crystal Structure of a Eukaryotic GEN1 Resolving Enzyme Bound to DNA. Cell Rep., 13, 2565–2575.

41. Liu, Y., Freeman, A., Déclais, A.-C., Gartner, A. and Lilley, D.M.J. (2018) Biochemical and Structural Properties of Fungal Holliday Junction-Resolving Enzymes. Methods Enzymol., 600, 543–568.

42. Laguri, C., Gilquin, B., Wolff, N., Romi-Lebrun, R., Courchay, K., Callebaut, I., Worman, H.J. and Zinn-Justin, S. (2001) Structural characterization of the LEM motif common to three human inner nuclear membrane proteins. Structure, 9, 503–511.

43. Lin, F., Blake, D.L., Callebaut, I., Skerjanc, I.S., Holmer, L., McBurney, M.W., Paulin-Levasseur, M. and Worman, H.J. (2000) MAN1, an inner nuclear membrane protein that shares the LEM domain with lamina-associated polypeptide 2 and emerin. J. Biol. Chem., 275, 4840–4847.

44. Shumaker, D.K., Lee, K.K., Tanhehco, Y.C., Craigie, R. and Wilson, K.L. (2001) LAP2 binds to BAF.DNA complexes: requirement for the LEM domain and modulation by variable regions. EMBO J., 20, 1754–1764.

45. Cai, M., Huang, Y., Suh, J.-Y., Louis, J.M., Ghirlando, R., Craigie, R. and Clore, G.M. (2007) Solution NMR structure of the barrier-to-autointegration factor-Emerin complex. J. Biol. Chem., 282, 14525–14535.

46. Cai, M., Huang, Y., Ghirlando, R., Wilson, K.L., Craigie, R. and Clore, G.M. (2001) Solution structure of the constant region of nuclear envelope protein LAP2 reveals two LEM-domain structures: one binds BAF and the other binds DNA. EMBO J., 20, 4399–4407.

47. Brachner, A. and Foisner, R. (2011) Evolvement of LEM proteins as chromatin tethers at the nuclear periphery. Biochem. Soc. Trans., 39, 1735–1741.

48. Matos, J. and West, S.C. (2014) Holliday junction resolution: regulation in space and time. DNA Repair, 19, 176–181.

49. Ip, S.C.Y., Rass, U., Blanco, M.G., Flynn, H.R., Skehel, J.M. and West, S.C. (2008) Identification of Holliday junction resolvases from humans and yeast. Nature, 456, 357– 361.

50. Chan, Y.W. and West, S. (2015) GEN1 promotes Holliday junction resolution by a coordinated nick and counter-nick mechanism. Nucleic Acids Res., 43, 10882–10892.

51. Carreira, R., Aguado, F.J., Hurtado-Nieves, V. and Blanco, M.G. (2022) Canonical and novel non-canonical activities of the Holliday junction resolvase Yen1. Nucleic Acids Res., 50, 259–280.

52. Gaillard, P.-H.L., Noguchi, E., Shanahan, P. and Russell, P. (2003) The endogenous Mus81-Eme1 complex resolves Holliday junctions by a nick and counternick mechanism. Mol. Cell, 12, 747–759.

53. Benitez, A., Sebald, M., Kanagaraj, R., Rodrigo-Brenni, M.C., Chan, Y.W., Liang, C.-C. and West, S.C. (2023) GEN1 promotes common fragile site expression. Cell Rep., 42, 112062.

54. Inagaki, H., Ohye, T., Kogo, H., Tsutsumi, M., Kato, T., Tong, M., Emanuel, B.S. and Kurahashi, H. (2013) Two sequential cleavage reactions on cruciform DNA structures cause palindrome-mediated chromosomal translocations. Nat. Commun., 4, 1592.

55. Zhao, J., Wang, G., Del Mundo, I.M., McKinney, J.A., Lu, X., Bacolla, A., Boulware, S.B., Zhang, C., Zhang, H., Ren, P., et al. (2018) Distinct Mechanisms of Nuclease-Directed DNA-Structure-Induced Genetic Instability in Cancer Genomes. Cell Rep., 22, 1200–1210.

56. Zlopasa, L., Brachner, A. and Foisner, R. (2016) Nucleo-cytoplasmic shuttling of the endonuclease ankyrin repeats and LEM domain-containing protein 1 (Ankle1) is mediated by canonical nuclear export- and nuclear import signals. BMC Cell Biol., 17, 23.

57. Dokshin, G.A., Ghanta, K.S., Piscopo, K.M. and Mello, C.C. (2018) Robust Genome Editing with Short Single-Stranded and Long, Partially Single-Stranded DNA Donors in Caenorhabditis elegans. Genetics, 210, 781–787.

58. Craig, A.L., Moser, S.C., Bailly, A.P. and Gartner, A. (2012) Methods for studying the DNA damage response in the Caenorhabdatis elegans germ line. Methods Cell Biol., 107, 321– 352.

59. Fitzgerald, D.J., Berger, P., Schaffitzel, C., Yamada, K., Richmond, T.J. and Berger, I. (2006) Protein complex expression by using multigene baculoviral vectors. Nat. Methods, 3, 1021– 1032.

60. Prazeres, D.M., Schluep, T. and Cooney, C. (1998) Preparative purification of supercoiled plasmid DNA using anion-exchange chromatography. J. Chromatogr. A, 806, 31–45.

61. Jumper, J., Evans, R., Pritzel, A., Green, T., Figurnov, M., Ronneberger, O., Tunyasuvunakool, K., Bates, R., Žídek, A., Potapenko, A., et al. (2021) Highly accurate protein structure prediction with AlphaFold. Nature, 596, 583–589.

62. Abramson, J., Adler, J., Dunger, J., Evans, R., Green, T., Pritzel, A., Ronneberger, O., Willmore, L., Ballard, A.J., Bambrick, J., et al. (2024) Accurate structure prediction of biomolecular interactions with AlphaFold 3. Nature, 630, 493–500.

