## Supplementary data for "Functional dissection of the conserved *C. elegans* LEM-3/ANKLE1 nuclease reveals a crucial requirement for the LEM-like and GIY-YIG domains for DNA bridges processing"

### Figure S1

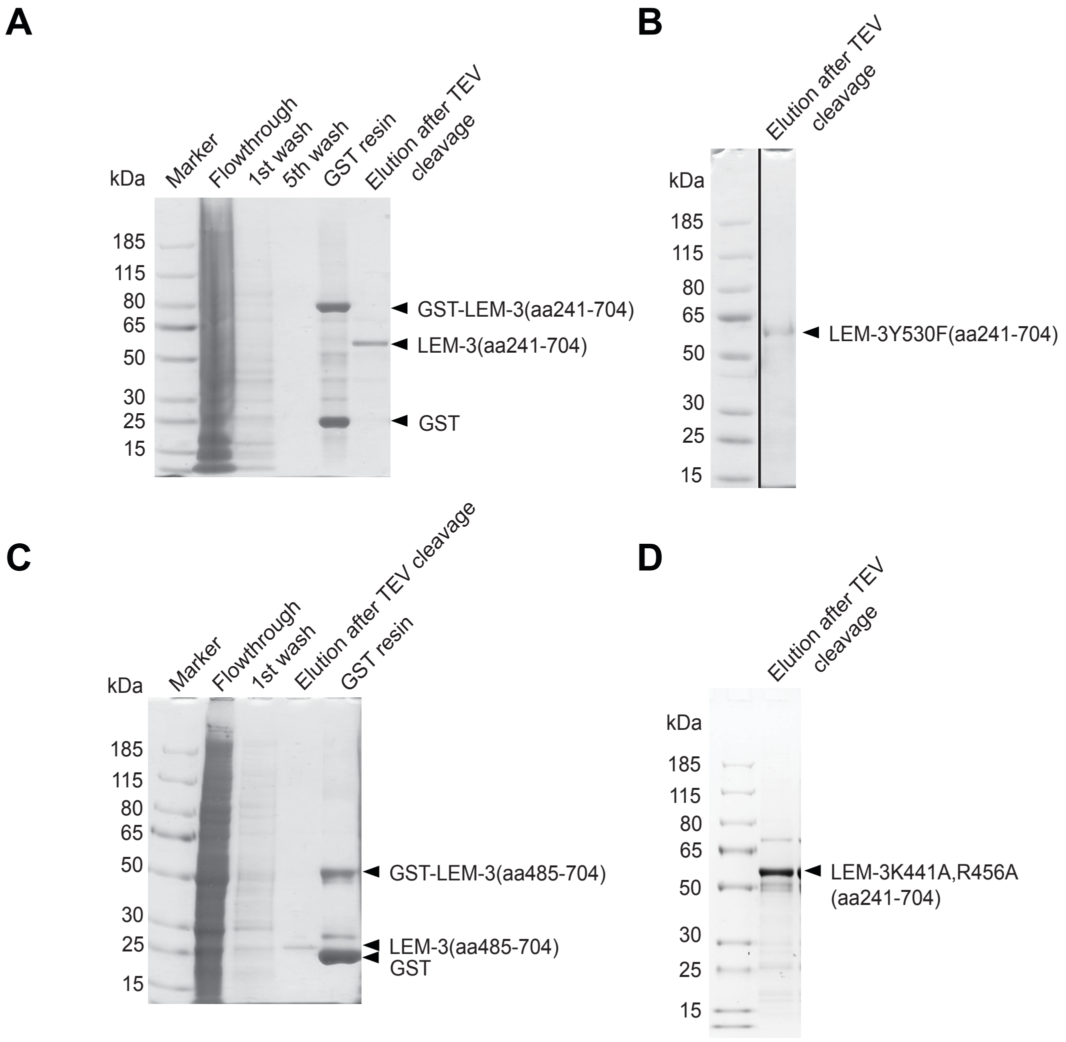
A
Marker
Flowthrough
1st wash
5th wash
GST resin
Elution after TEV cleavage
kDa
185
115
80
65
50
30
25
15
GST-LEM-3(aa241-704)
LEM-3(aa241-704)
GST
B
Elution after TEV cleavage
kDa
185
115
80
65
50
30
25
15
LEM-3Y530F(aa241-704)
C
Marker
Flowthrough
1st wash
Elution after TEV cleavage
GST resin
kDa
185
115
80
65
50
30
25
15
GST-LEM-3(aa485-704)
LEM-3(aa485-704)
GST
D
Elution after TEV cleavage
kDa
185
115
80
65
50
30
25
15
LEM-3K441A,R456A
(aa241-704)

**Figure S1: Purification of the different LEM-3 derivates used in this study using the baculovirus-based expression system (related to Figure 1, 3, 4 and 5).** Flowthrough, 1st wash, 5th wash, GST resin, and Elution after TEV cleavage are indicated **(A)** LEM-3(241-704), **(B)** LEM-3Y530F(241-704), **(C)** LEM-3(485-704) and **(D)** LEM-3K441A;R456A(241-704). Uncropped versions of panels B and D are shown in Supplementary Figures S11A and B, respectively.

Figure S2

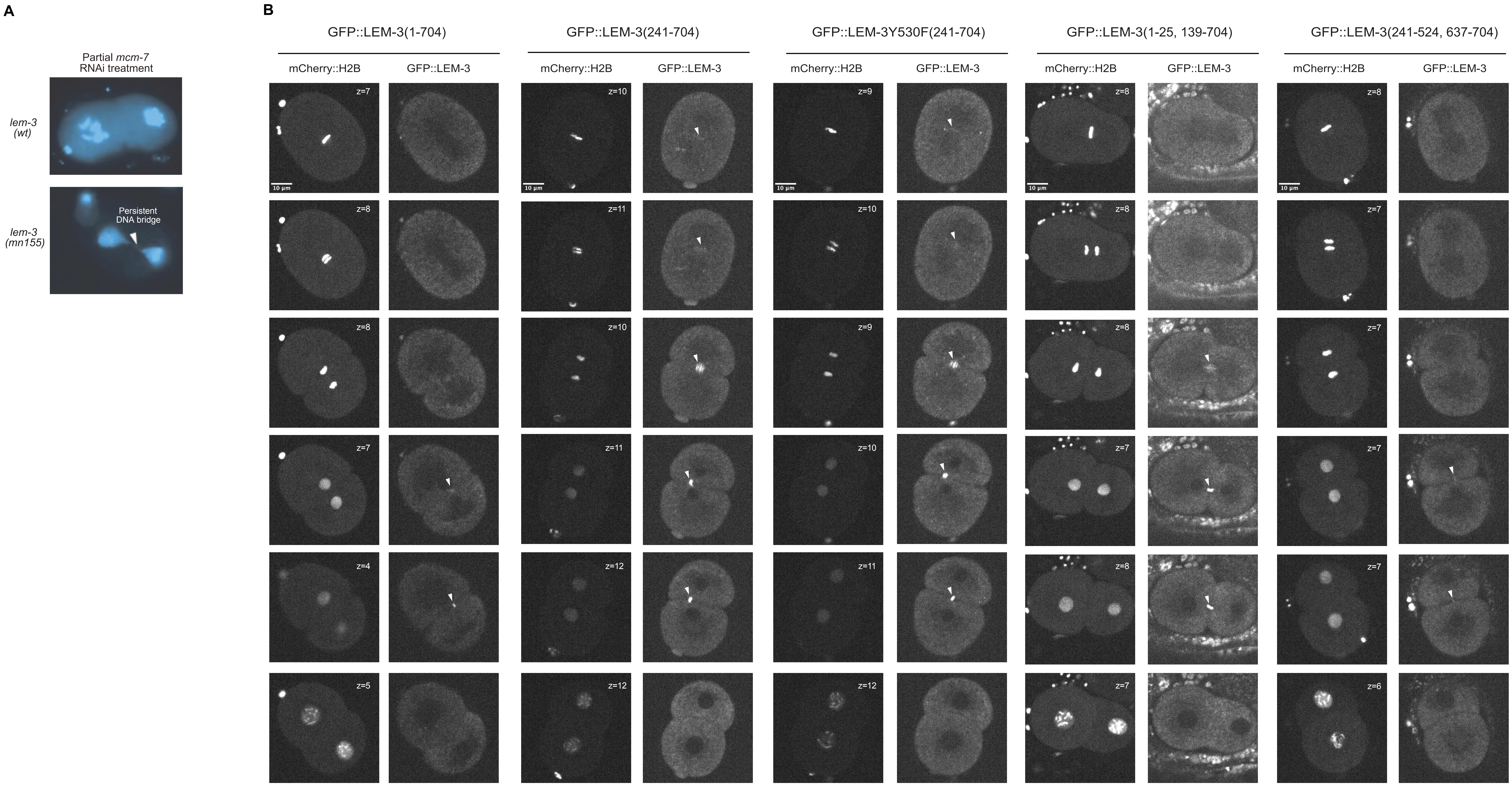
A
Partial mcm-7
RNAi treatment
lem-3
(wt)
lem-3
(mn155)
Persistent
DNA bridge
B
GFP::LEM-3(1-704)
GFP::LEM-3(241-704)
GFP::LEM-3Y530F(241-704)
GFP::LEM-3(1-25, 139-704)
GFP::LEM-3(241-524, 637-704)
mCherry::H2B
GFP::LEM-3
z=7
z=8
z=8
z=7
z=4
z=5
z=10
z=11
z=10
z=11
z=12
z=12
z=9
z=10
z=9
z=10
z=11
z=12
z=8
z=8
z=8
z=7
z=8
z=7
z=8
z=7
z=7
z=7
z=7
z=6
10 µm

**Figure S2: Partial *mcm-7* RNAi experiment and time-lapse imaging of the first cell division of various GFP::LEM-3 mutant strains in the absence of DNA bridges (related to Figure 1). (A)** Upon partial *mcm-7* RNAi, persistent DNA bridges are revealed by DAPI staining. **(B)** mCherry:H2B and GFP channels are shown. The time (post-anaphase onset) and the Z-position in the Z-stack for each snapshot are indicated. For all the GFP::LEM-3 mutant strains and the wild-type GFP::LEM-3 strain, the LUT (95-150) was used.

### Figure S3

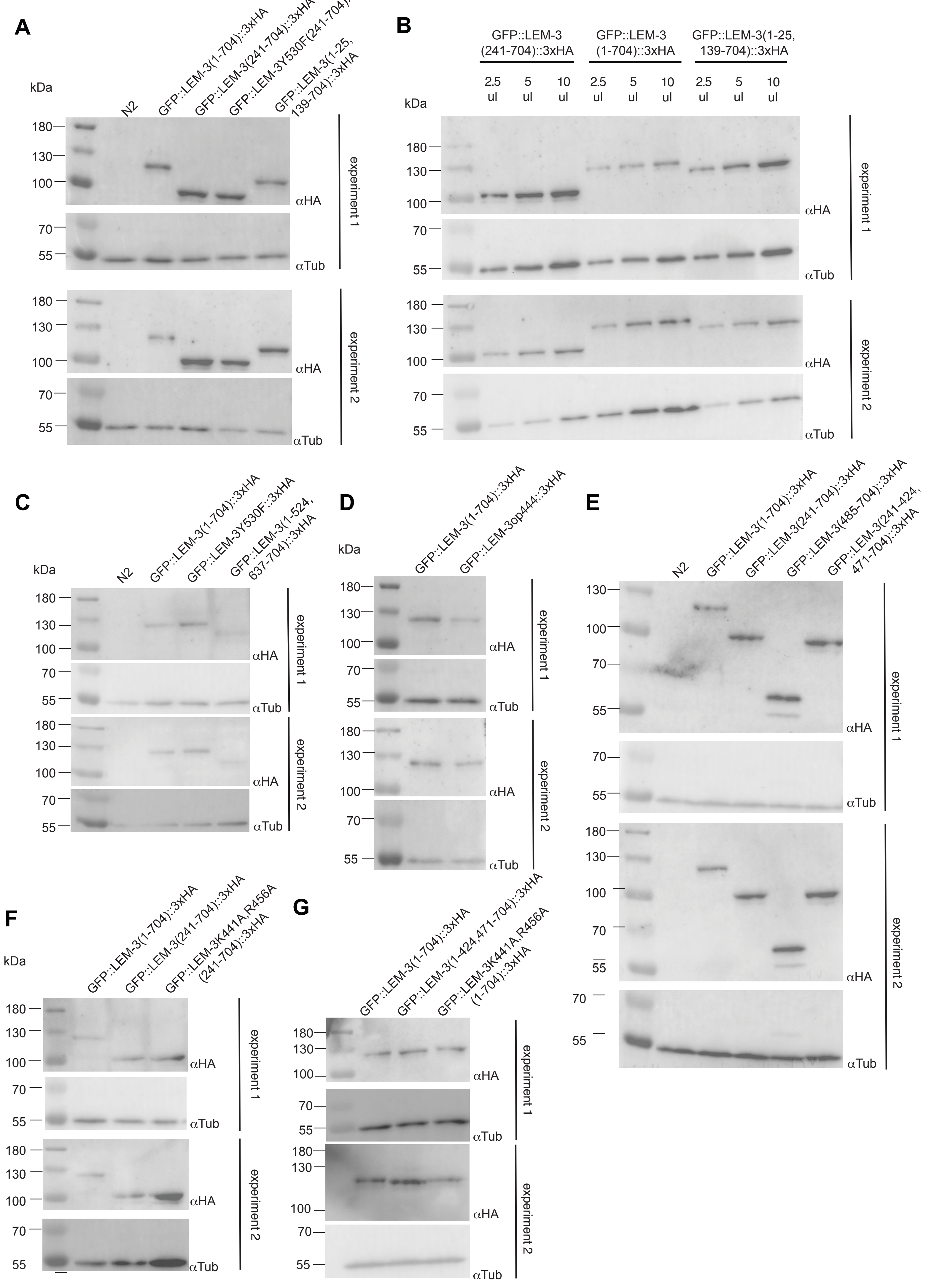
A
kDa
N2
GFP::LEM-3(1-704)::3xHA
GFP::LEM-3(241-704)::3xHA
GFP::LEM-3Y530F(241-704)::3xHA
GFP::LEM-3(1-25, 139-704)::3xHA
180
130
100
70
55
αHA
αTub
experiment 1
experiment 2
B
GFP::LEM-3 (241-704)::3xHA
GFP::LEM-3 (1-704)::3xHA
GFP::LEM-3(1-25, 139-704)::3xHA
2.5 ul
5 ul
10 ul
C
GFP::LEM-3Y530F::3xHA
GFP::LEM-3(1-524, 637-704)::3xHA
D
GFP::LEM-3op444::3xHA
E
GFP::LEM-3(485-704)::3xHA
GFP::LEM-3(241-424, 471-704)::3xHA
F
GFP::LEM-3K441A,R456A (241-704)::3xHA
G
GFP::LEM-3(1-424,471-704)::3xHA
GFP::LEM-3K441A,R456A (1-704)::3xHA

**Figure S3: Western-blot analysis of the level of the different GFP::LEM-3 derivates (Related to Figures 1, 2, 4, 5, and 6).** As described in the methods, the different strains were grown at 25°C for 3 days. Total proteins were extracted and analyzed by SDS-PAGE and Western using anti-HA and anti-Tubulin antibodies. For each panel, two independent experiments were performed.

### Figure S4

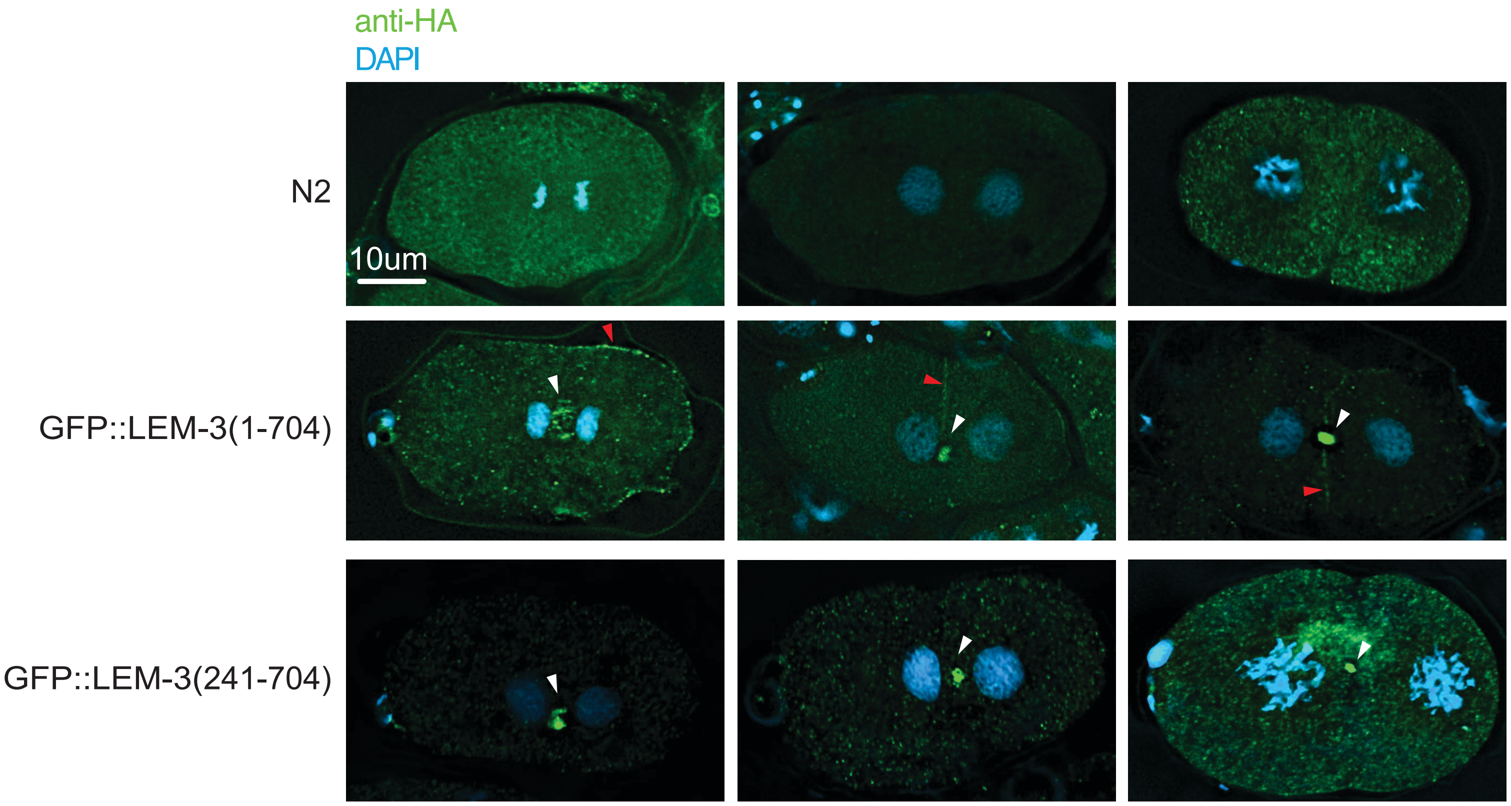

**Figure S4: Immunostaining of GFP::LEM-3(1-704) and GFP::LEM-3(241-704) (related to Figure 1).** As described in the methods, embryos were subjected to the ‘freeze-crack’ method and stained with anti-GFP antibodies. Early stages of the first cell division were imaged by epifluorescence. Deconvolved images are shown. N2 embryos were used as a control. White arrowheads indicate the LEM-3 foci at the midzone and midbody while red arrowheads indicate LEM-3 at the plasma membrane.

### Figure S5

**A**

```
                      Y530                 Y556 K559             R564 H568
LEM-3    525  YNAFCYLIMDP 535   553 SIFYVGKGSK 562   561 SKNRPLAHFIDARNERRD 578
ANKLE1   449  KSSFTYLLLDP 459   484 AIFYVGKGTR 492   491 TRARPYVHLWEALGHHG 507
Cg-SLX1    9  DFYGCYLLQSI 19    23  QSFYIGSTPN 32    32 NPVRRLRQHNGS 44
T4endoII 32   KYNVIYAIAIN 42    44  ELVYIGKTKN 53    54  LRKRINYYRTAI 65
UvrC      14  EEPGVYIFKNK 24    26  VPIYIGKAKR 35    35 RLSNRLRSYLNP 46
Hpy188I  58   NEGLVYVFVIQ 68    70  KIFKIGHSIT 79    81  ITKRVQSYNCG 91
                β sheet1             β sheet2              α helix1

                                            E620
LEM-3    602 IPRHEISHGV  611   613 DEEAFVREASIIEAVKLKNLRNKKAGEFH 641
ANKLE1   534 VSLHCFQHVV  543   539 FQHVVAVEAYTREACIVEALGIQTLTNQK 567
Cg-SLX1   60 WEMVAIVYGF  69    71  SRIAALQFEHAWQHGYQTRYIKSQDRVVKT 100
T4endoII 90  VEFYARQCFN  99    112  ATIDLEEPLFIKLFNPPWNI 131
UvrC      59 DELETIVVMN  68    68  NREAFILEANLIKKYRPKYNV 89
Hpy188I  119 VQVYAFFPEQ  128   143  STSKRAENVILENFIKNHNKKPIGCT 168
              β sheet3                  α helix2
```

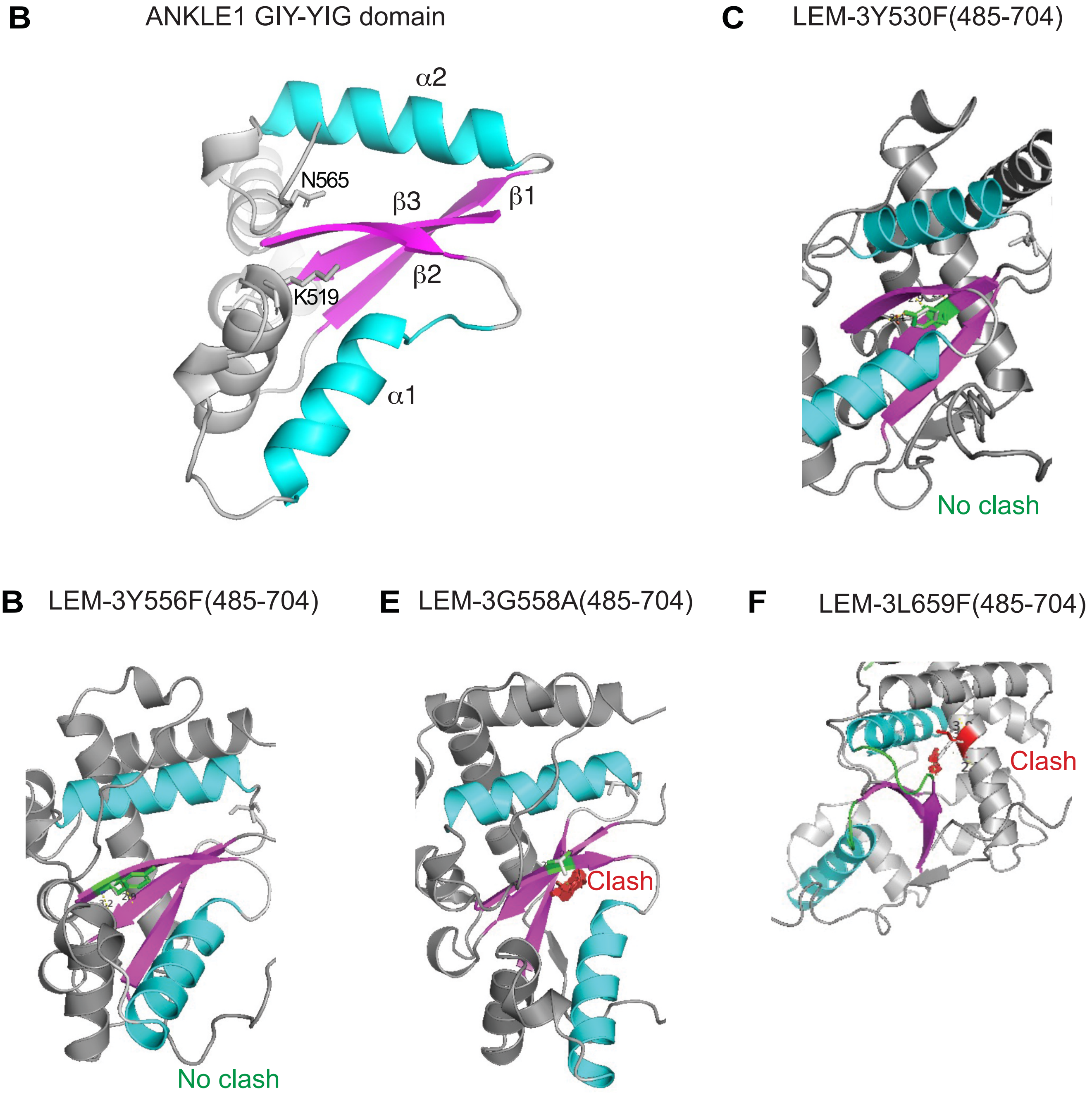

**Figure S5: Conservation of the LEM-3 GIY-YIG domain and predicted structure of various LEM-3 mutants (related to Figure 2). (A)** Sequence alignment of the GIY-YIG domain of different GIY-YIG nucleases **(B)** Predicted structure by AlphaFold of ANKLE1 GIY-YIG domain with the position of the K519 and N565 residues. **(C-F)** Predicted structure by AlphaFold of LEM-3Y530F(485-704), LEM-3Y556F(485-704), LEM-3G558A(485-704) and LEM-3L659F(485-704). Predicted structural clashes are indicated.

### Figure S6

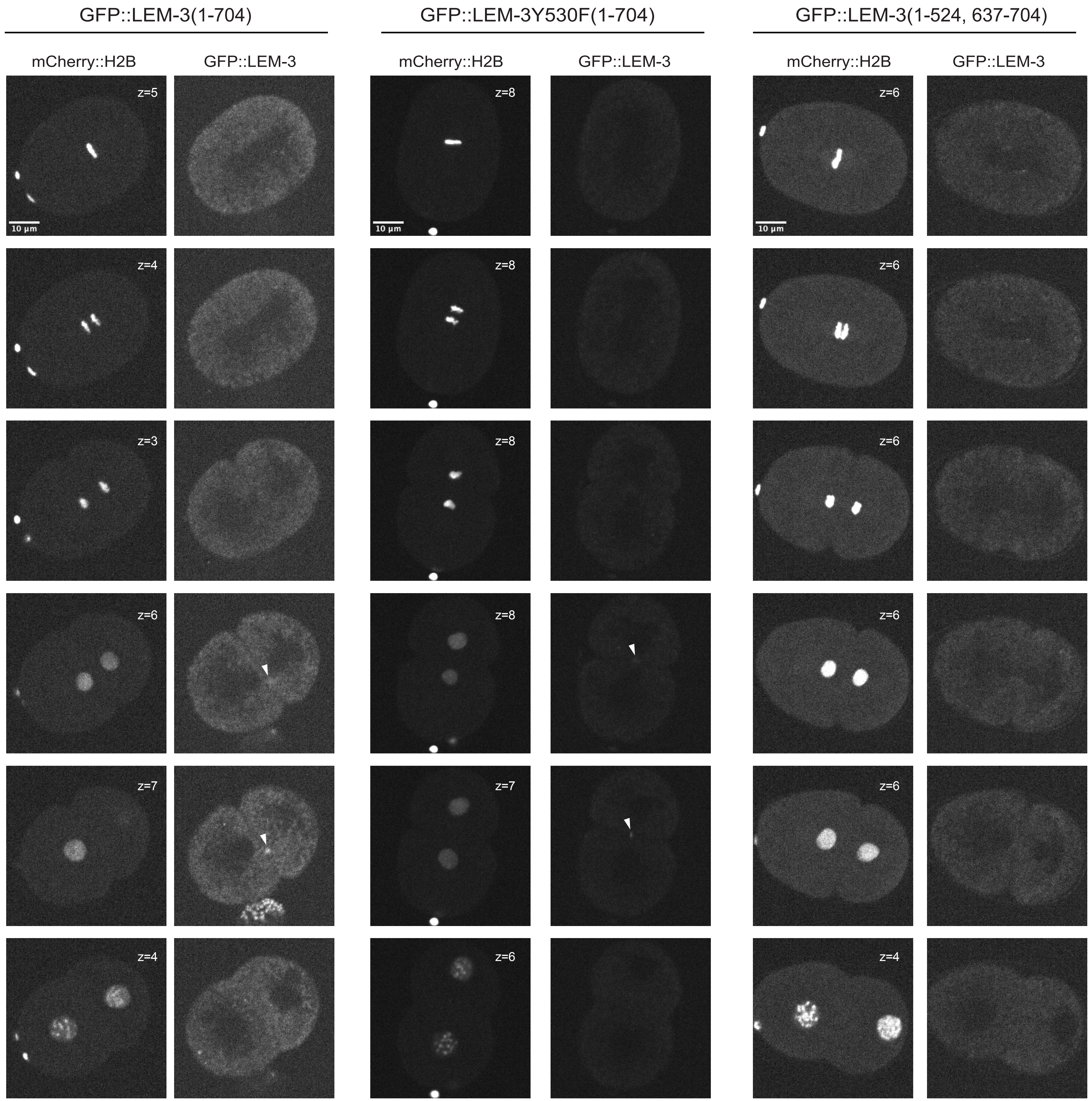

GFP::LEM-3(1-704)
GFP::LEM-3Y530F(1-704)
GFP::LEM-3(1-524, 637-704)
mCherry::H2B
GFP::LEM-3
z=5
z=4
z=3
z=6
z=7
z=8
10 µm

**Figure S6: Time-lapse imaging of the first cell division of various GFP::LEM-3 mutant strains in the absence of DNA bridges (related to Figure 2).** mCherry:H2B and GFP channels are shown. The time (post-anaphase onset) and the Z-position in the Z-stack for each snapshot are indicated. For all the GFP::LEM-3 mutant strains and the wild-type GFP::LEM-3 strain, the LUT (95-150) was used.

#### Figure S7

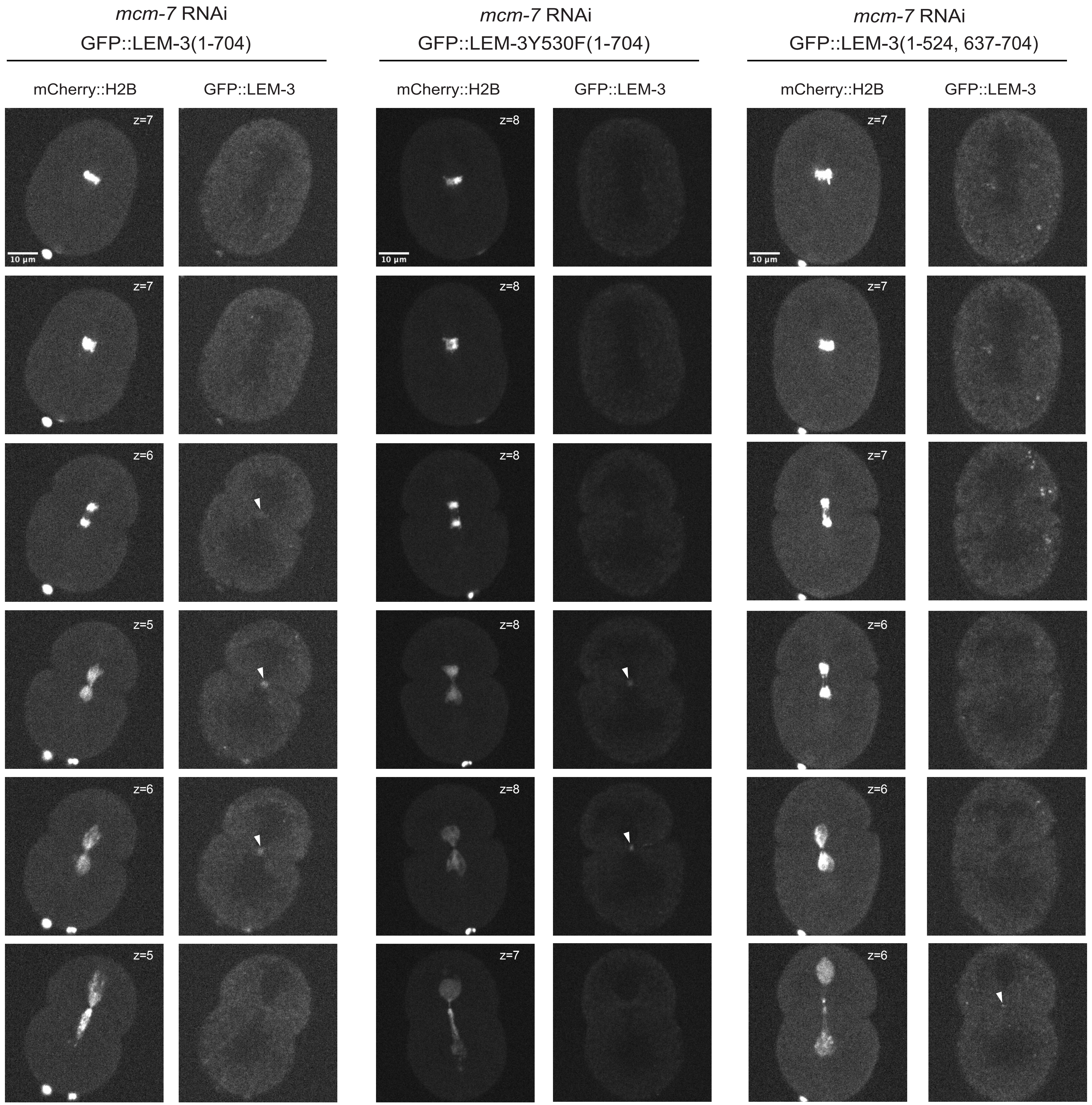
*mcm-7* RNAi
GFP::LEM-3(1-704)
*mcm-7* RNAi
GFP::LEM-3Y530F(1-704)
*mcm-7* RNAi
GFP::LEM-3(1-524, 637-704)
mCherry::H2B
GFP::LEM-3
mCherry::H2B
GFP::LEM-3
mCherry::H2B
GFP::LEM-3
z=7
z=8
z=7
10 µm
z=7
z=8
z=7
z=6
z=8
z=7
z=5
z=8
z=6
z=6
z=8
z=6
z=5
z=7
z=6

**Figure S7: Time-lapse imaging of the first cell division of various GFP::LEM-3 mutant strains in the presence of DNA bridges (related to Figure 2).** mCherry:H2B and GFP channels are shown. The time (post-anaphase onset) and the Z-position in the Z-stack for each snapshot are indicated. For all the GFP::LEM-3 mutant strains and the wild-type GFP::LEM-3 strain, the LUT (95-150) was used.

### Figure S8

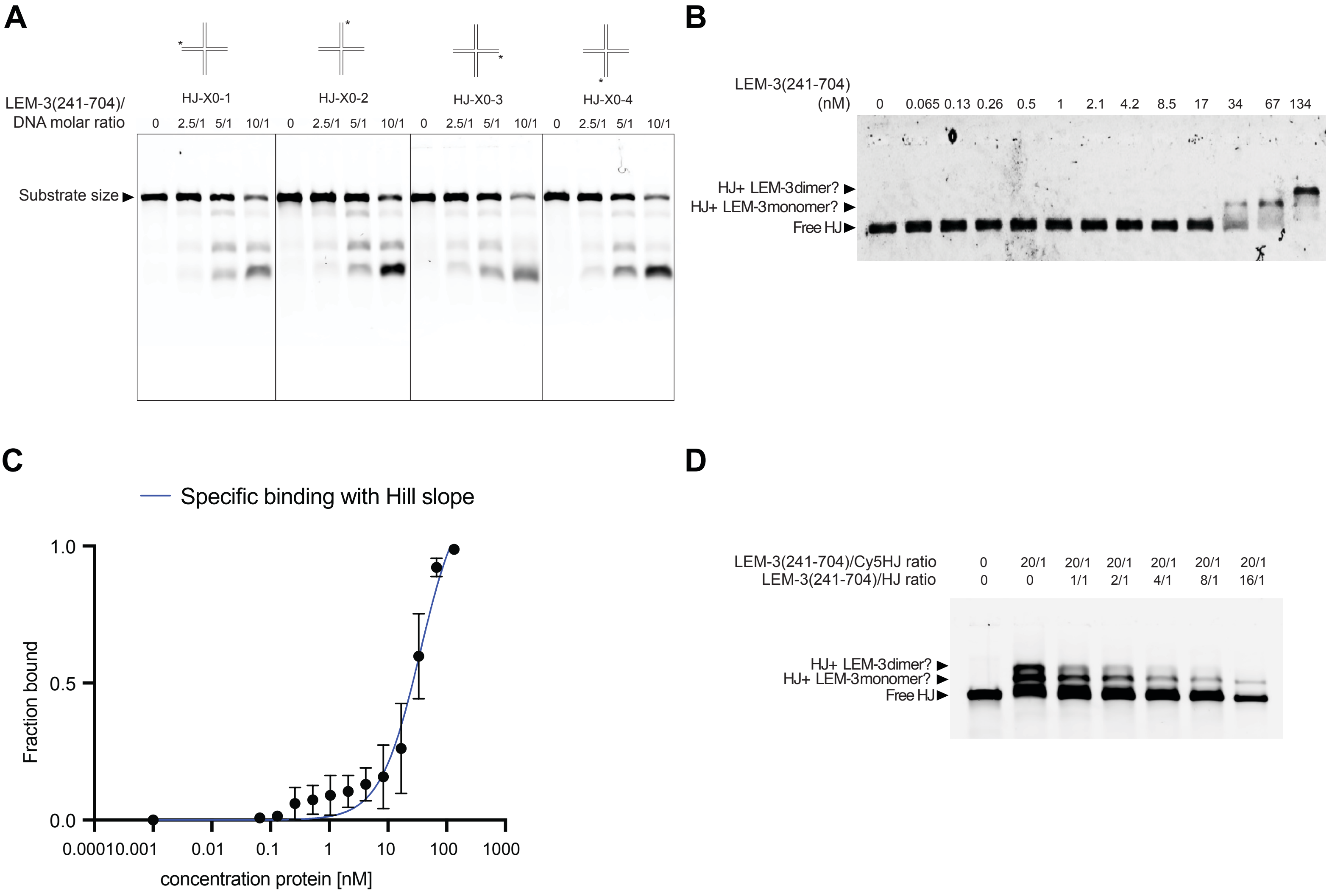
A
HJ-X0-1
HJ-X0-2
HJ-X0-3
HJ-X0-4
LEM-3(241-704)/
DNA molar ratio
0 2.5/1 5/1 10/1 0 2.5/1 5/1 10/1 0 2.5/1 5/1 10/1 0 2.5/1 5/1 10/1
Substrate size
B
LEM-3(241-704)
(nM)
0 0.065 0.13 0.26 0.5 1 2.1 4.2 8.5 17 34 67 134
HJ+ LEM-3dimer?
HJ+ LEM-3monomer?
Free HJ
C
Specific binding with Hill slope
Fraction bound
1.0
0.5
0.0
0.00010.001 0.01 0.1 1 10 100 1000
concentration protein [nM]
D
LEM-3(241-704)/Cy5HJ ratio 0 20/1 20/1 20/1 20/1 20/1 20/1
LEM-3(241-704)/HJ ratio 0 0 1/1 2/1 4/1 8/1 16/1
HJ+ LEM-3dimer?
HJ+ LEM-3monomer?
Free HJ

**Figure S8: Mapping of the cleavage site of HJ by LEM-3 and measure of the affinity of LEM-3 for HJ (related to Figure 3). (A)** Nuclease activity of LEM-3(241-704) on HJ, where the labeling was performed on each of the 4 strands. Different protein/DNA substrate molar ratios were used, and incubation was performed at 37 C for 10 min. The cleavage products were analyzed by 6% neutral PAGE. The size of the DNA substrates is indicated. **(B)** Various concentrations of LEM-3 protein were incubated with 0.13nM HJ-X0-1, and the resulting products were analyzed on a 6% neutral PAGE. **(C)** The HJ bound fraction was plotted as a function of LEM-3(241-704) concentration and the data was fitted to the Hill equation**. (D)** Labeled HJ-X0-1 (Cy5HJ) was incubated with LEM-3(241-704) and the ratio of shifted protein was measured by gel shift assay in the presence of increasing amounts of unlabeled HJ substrate.

### Figure S9

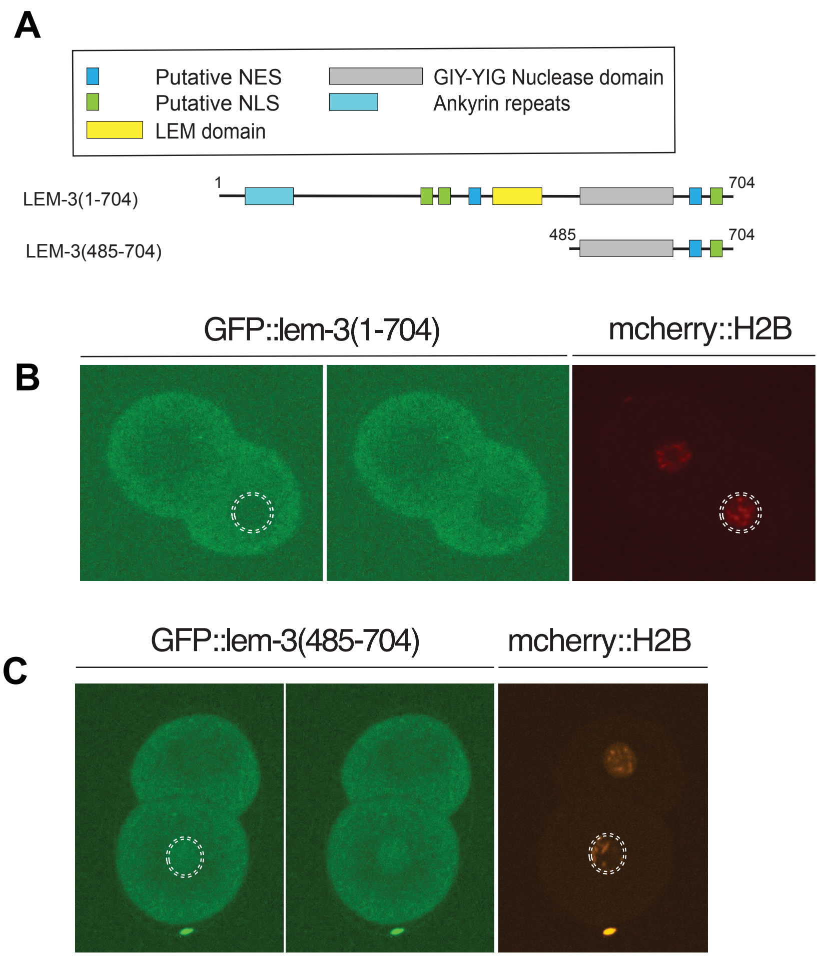
A
Putative NES
Putative NLS
LEM domain
GIY-YIG Nuclease domain
Ankyrin repeats
LEM-3(1-704)
1
704
LEM-3(485-704)
485
704
GFP::lem-3(1-704)
mcherry::H2B
B
GFP::lem-3(485-704)
mcherry::H2B
C

**Figure S9: LEM-3(485-704) derivate partially mis-localized to the nucleus. (A)** Schematic of the different LEM-3 derivates used in this figure. **(B)** GFP::LEM-3(1-704) is excluded from the nucleus (dotted circle). **(C)** GFP:LEM-3 (485-704) is not excluded from the nucleus (dotted circle).

### Figure S10

**A** Alignment of LEM domain in ANKLE1 family

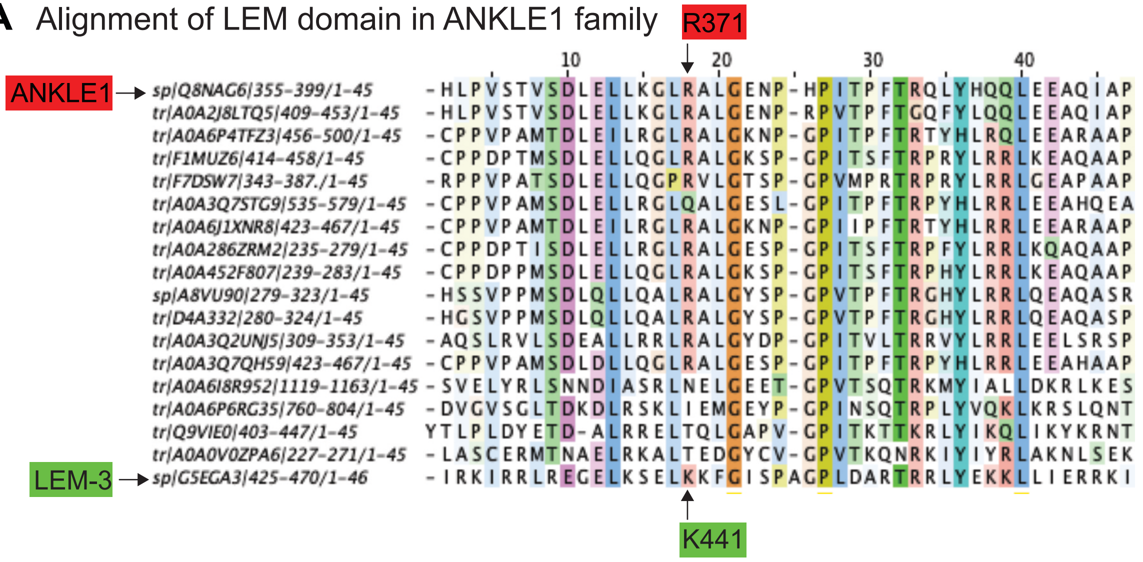


**B** Alignment of LEM domain in ANKLE1 family to LEM like domain in LAP2 family

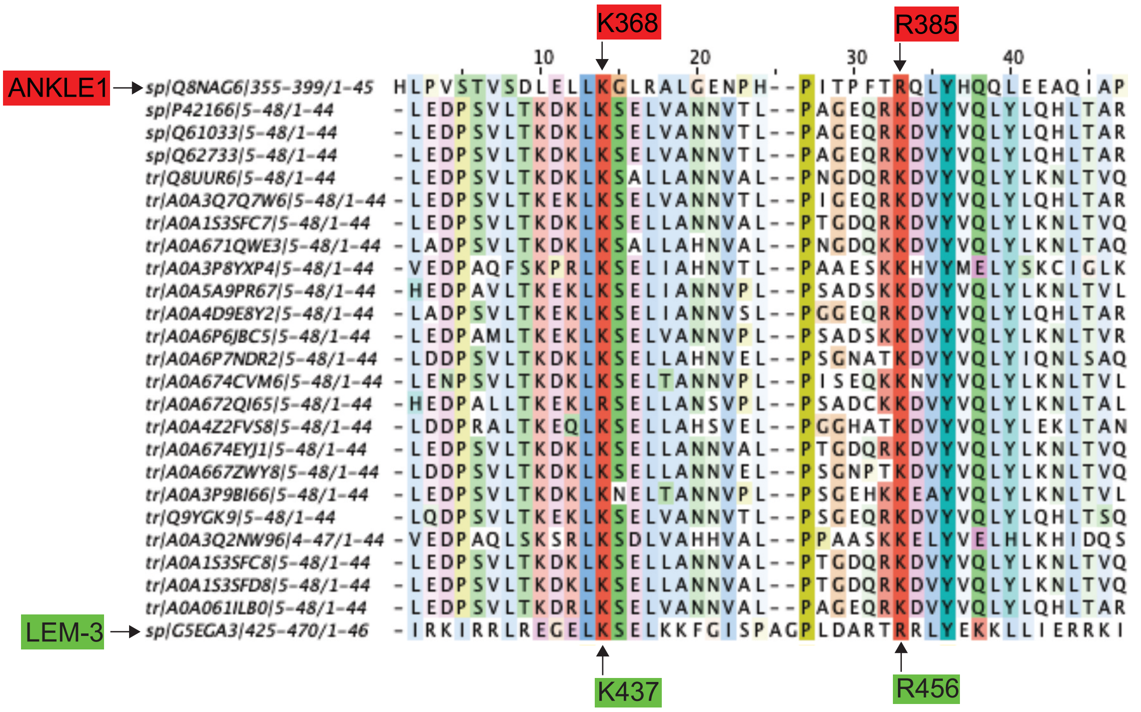


**C** Alignment of LEM domain in ANKLE1 family to LEM domain in LAP2 family

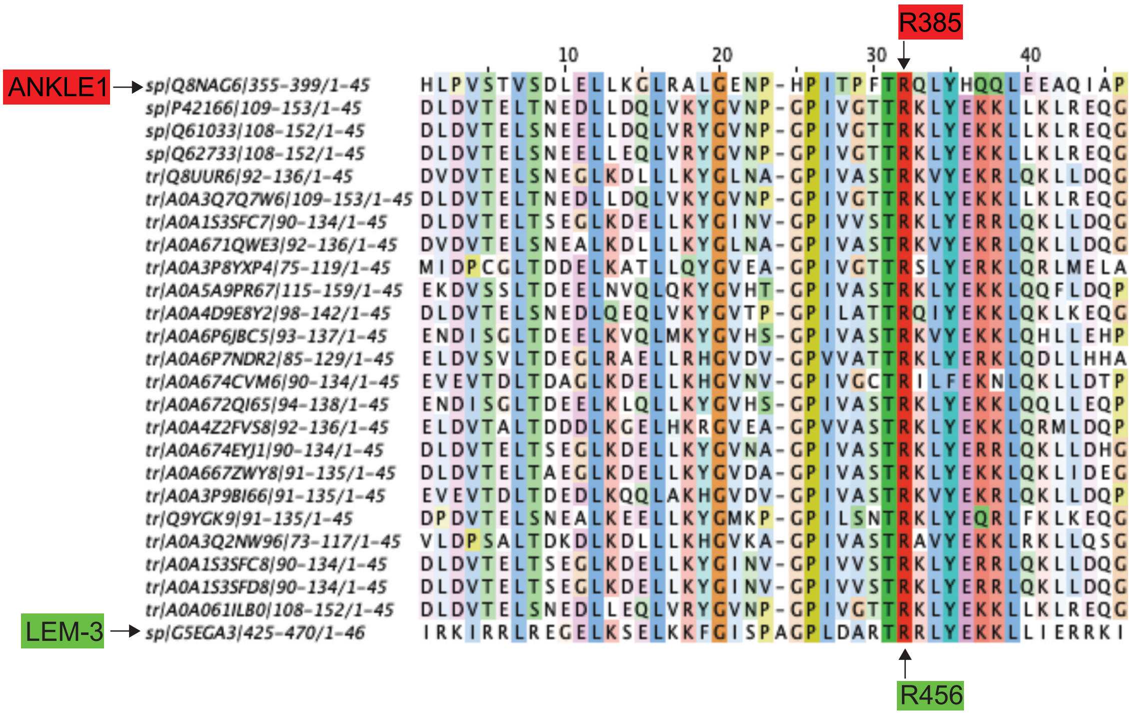

**Figure S10: Sequence alignments of LEM and LEM-like domains. (A)** Alignment of LEM domain in ANKLE1 family. **(B)** Alignment of LEM domain in ANKLE1 family to LEM-like domain in LAP2 family. **(C)** Alignment of LEM domain in ANKLE1 family to LEM domain in LAP2 family.

### Figure S11

A

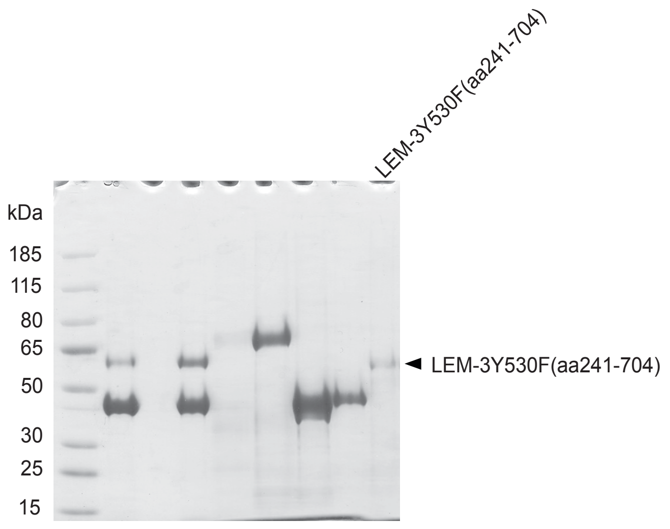


B

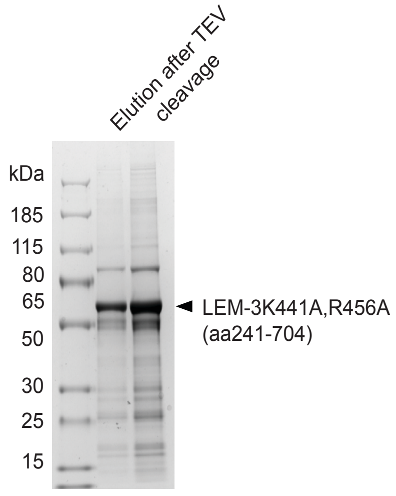

**Figure S11: Uncropped image of coomassie gel. (A)** uncropped image of Figure S1B **(B)** uncropped image of Figure S1D.

**Table S1: List of strains used in this study**

| Strain name | Genotype | Source |
|---|---|---|
| N2 | Wild-type | CGC |
| TG4397 | *lem-3(cop859 [Plem-3:GFP::Stag::lem-3(1-704):3'UTRlem-3]) I ; odIs57 [Ppie-1:mcherry::H2B]* | Gartner lab |
| SP483 | *lem-3 (mn155) I (premature stop codon at R190)* | CGC |
| TG4318 | *lem-3(op444) I* | Ye Hong |
| PHX1970 | *lem-3(syb1970 [Plem-3:GFP::Stag::lem-3E620A(1-704):3'UTRlem-3]) I ; odIs57 [Ppie-1:mcherry::H2B].*<br>*Generated in the background of TG4397 lem-3(cop859).* | SunyBiotech |
| PHX2062 | *lem-3(syb2062 [Plem-3:GFP::Stag::lem-3Y556F(1-704):3'UTRlem-3]) I ; odIs57 [Ppie-1:mcherry::H2B].*<br>*Generated in the background of TG4397 lem-3(cop859).* | SunyBiotech |
| PHX1949 | *lem-3(syb1949 [Plem-3:GFP::Stag::lem-3Y530F(1-704):3'UTRlem-3]) I ; odIs57 [Ppie-1:mcherry::H2B].*<br>*Generated in the background of TG4397 lem-3(cop859).* | SunyBiotech |
| PHX4324 | *lem-3(syb4324 [Plem-3:GFP::Stag::lem-3(1-524,637-704):3'UTRlem-3]) I ; odIs57 [Ppie-1:mcherry::H2B].*<br>*Generated in the background of TG4397 lem-3(cop859).* | SunyBiotech |
| PHX3292 | *lem-3(syb3292 [Plem-3:GFP::Stag::lem-3(485-704):3'UTRlem-3]) I ; odIs57 [Ppie-1:mcherry::H2B].*<br>*Generated in the background of TG4397 lem-3(cop859).* | SunyBiotech |
| PHX1935 | *lem-3(syb1935 [Plem-3:GFP::Stag::lem-3(1-424,471-704):3'UTRlem-3]) I ; odIs57 [Ppie-1:mcherry::H2B].*<br>*Generated in the background of TG4397 lem-3(cop859).* | SunyBiotech |
| PHX2728 | *lem-3(syb2728 [Plem-3:GFP::Stag::lem-3(241-704):3'UTRlem-3]) I ; odIs57 [Ppie-1:mcherry::H2B].*<br>*Generated in the background of TG4397 lem-3(cop859).* | SunyBiotech |
| PHX7187 | *lem-3(syb7187[Plem-3::GFP::Stag::lem-3Y530F(485-704):3'UTRlem-3]) I ; odIs57 [Ppie-1:mcherry::H2B].*<br>*Generated in the background of TG4397 lem-3(syb3292).* | SunyBiotech |
| PHX6759 | *lem-3(syb6759{full KO]) I.*<br>*Generated in the background of N2(wt).* | SunyBiotech |
| TG4884 | *lem-3(gt3440[Plem-3::GFP::Stag::lem-3Y530F(241-704):3'UTRlem- 3]) I ; odIs57 [Ppie-1:mcherry::H2B].*<br>*Generated in the background of TG4397 lem-3(syb2728).* | Stéphane Rolland |
| TG4887 | *lem-3(gt3443[Plem-3::GFP::Stag::lem-3K441A;R456A(241-704):3'UTRlem-3]) I ;odIs57[Ppie1:mcherry::H2B].* | Stéphane Rolland |

| | | |
|---|---|---|
| | *Generated in the background of TG4861 lem-3(gt3420).* | |
| TG4892 | *lem-3(gt3445[Plem-3::GFP::Stag::lem-3(241-424,471-704):3'UTRlem-3] )I ;odIs57[Ppie1:mcherry::H2B].*<br>*Generated in the background of TG4397 lem-3(syb2728).* | Stéphane Rolland |
| PHX7665 | *lem-3(syb7665[Plem-3::GFP::Stag::lem-3(1-25,139-704):3'UTRlem-3] ) I ; odIs57 [Ppie-1:mcherry::H2B].*<br>*Generated in the background of TG4397 lem-3(cop859).* | SunyBiotech |
| TG4688 | *lem-3(gt3382[Plem-3::eGFP::Stag::lem-3(K441A;R456A)]) I; odIs57 [Ppie-1:mcherry::H2B].*<br>*Generated in the background of TG4397 lem-3(cop859).* | Stéphane Rolland |
| PHX7973 | *lem-3(syb7973[Plem-3::GFP::Stag::lem-3(1-424):3'UTRlem-3] ) I ; odIs57 [Ppie-1:mcherry::H2B].*<br>*Generated in the background of TG4397 lem-3(cop859).* | SunyBiotech |
| TG4584 | *lem-3(gt3329[Plem-3::GFP::Stag::lem-3(1-704)::3xHA:3'UTRlem-3] ) I ; odIs57 [Ppie-1:mcherry::H2B].*<br>*Generated in the background of TG4397 lem-3(cop859).* | Stéphane Rolland |
| TG4894 | *lem-3(gt3447[Plem-3::GFP::Stag::lem-3(241-704)::3xHA:3'UTRlem-3] ) I ; odIs57 [Ppie-1:mcherry::H2B].*<br>*Generated in the background of PHX2728 lem-3(syb2728).* | Stéphane Rolland |
| TG4969 | *lem-3(gt3500[Plem-3::GFP::Stag::lem-3Y530F(241-704)::3xHA:3'UTRlem-3] ) I ; odIs57 [Ppie-1:mcherry::H2B].*<br>*Generated in the background of TG4884 lem-3(gt3440).* | Stéphane Rolland |
| TG4968 | *lem-3(gt3499[Plem-3::GFP::Stag::lem-3(1-25,139-704)::3xHA:3'UTRlem-3] ) I ; odIs57 [Ppie-1:mcherry::H2B].*<br>*Generated in the background of PHX7665 lem-3(syb7665).* | Stéphane Rolland |
| TG4828 | *lem-3(gt3403[Plem-3::GFP::Stag::lem-3Y530F(1-704)::3xHA:3'UTRlem-3] ) I ; odIs57 [Ppie-1:mcherry::H2B].*<br>*Generated in the background of PHX1949 lem-3(syb1949)* | Stéphane Rolland |
| TG4896 | *lem-3(gt3449 [Plem-3:GFP::Stag::lem-3(1-524,637-704)::3xHA:3'UTRlem- 3]) I ; odIs57 [Ppie-1:mcherry::H2B].*<br>*Generated in the background of PHX4324 lem-3(syb4324).* | Stéphane Rolland |
| TG4897 | *lem-3(gt3450 [Plem-3:GFP::Stag::lem-3(485-704)::3xHA:3'UTRlem- 3]) I ; odIs57 [Ppie-1:mcherry::H2B].*<br>*Generated in the background of PHX3292 lem-3(syb3292).* | Stéphane Rolland |
| TG4951 | *lem-3(gt3489[Plem-3::GFP::Stag::lem-3(241-424,471-704)::3xHA:3'UTRlem- 3] ) I ;odIs57[Ppie1:mcherry::H2B].*<br>*Generated in the background of TG4892 lem-3(gt3445).* | Stéphane Rolland |
| TG4952 | *lem-3(gt3491[Plem-3::GFP::Stag::lem-3K441A;R456A(241-704)::3xHA :3'UTRlem- 3] ) I ;odIs57[Ppie1:mcherry::H2B].*<br>*Generated in the background of TG4887 lem-3(gt3443).* | Stéphane Rolland |
| TG4965 | *lem-3(gt3497 [Plem-3:GFP::Stag::lem-3(L659F)::3xHA:3'UTRlem-3]) I ; odIs57 [Ppie-1:mcherry::H2B].* | Stéphane Rolland |

| | | |
|---|---|---|
| | *Generated in the background of TG4553 lem-3(gt3326)* | |
| TG4639 | *lem-3(gt3364[Plem-3::eGFP::Stag::lem-3(deltaLEM)::3xHA]) I ; odIs57 [Ppie-1:mcherry::H2B].*<br>*Generated in the background of PHX1935 lem-3(syb1935)* | Stéphane Rolland |
| TG4711 | *lem-3(gt3396[Plem-3::eGFP::lem-3(K441A;R456A)::3HA]) I ; odIs57 [Ppie-1:mcherry::H2B].*<br>*Generated in the background of TG4688 lem-3(gt3382)* | Stéphane Rolland |

**Table S2.** crRNA and ssODN used for genome editing

| | name | sequence | Used to generate the following strains |
|---|---|---|---|
| crRNA | crRNAlem-3op444 | ATGGAACATTTTTGCTGGAT**AGG** | TG4553 |
| | crRNAlem-3-Y530F | TCGAATTTTGGGAAGCAATG**TGG** | TG4884 |
| | crRNAlem-3-2 | CTTCTTGTACGAGCATCCAG**AGG** | TG4861, TG4887 |
| | crRNAlem-3-1 | AAGAGCGAGCTGAAAAAGTT**CGG** | TG4887 |
| | crRNAlem-3L | ACACCGACCACAGTCGATGA**TGG** | TG4892 |
| | crRNAlem-3R | TACGAACCGTGGATATTCAC**CGG** | TG4892 |
| | cRNA-lem3Cterm | GACGTGGAGCAGGTGGTGGA**CGG** | TG4584, TG4894, TG4969, TG4968, TG4828, TG4896, TG4897, TG4951, TG4953, TG4965 |
| ssODN | ssODNlem-3op444 | GCTGGGACAATATCACTAAATCAGAATATGGAAC ATTTTTTCTCGACAGgtgaacatctcgccattatttttcaatctaaa | TG4553 |
| | ssODNlem-3-Y530F | GTGGAAATGGATTCGGATATAATGCGTTTTGCTTC CTCATTATGGATCCTCGAATTCTGGGAAGCAACGT GGAGAACCTTACACTTGAAACCTTTGTACGATCA | TG4884 |
| | ssODN-R456A | AGCGAGCTGAAAAAGTTCGGAATCTCTCCAGCAG GACCTCTGGACGCTCGTACAGCTAGACTATATGA GAAGAAACTCCTGATTGAAAGACGGAAAATT | TG4861 |
| | ssODN-lem-3-triple | GGAGAAATCAGAAAAATTCGACGT CTTCGAGAAG GAGAACTGGCTAGCGAGCTGGCTAAGTTCGGAAT CTCTCCAGCAGGACCTCTGGACGCTCGTACAGCT AGACTATATGAGAAGAAACTCCTGATTGAAAGAC GGAAAATT | TG4887 |
| | ssODNdeltaLEM | TCATCTGCGGAAGACGACAAAGAAGCAGAAGTAT CAACACCGACCACAGTTGACGATGGAGAAACGAA CCGTGGATACTCGCCGGATGCTGACGTTGTTTCA TGTGTAAGTTATATATTTTCATTTCTC | TG4892 |
| | ssODNlem-3HA | TTTATCCATATGTGAACAATCGACGTGGAGCAGG TGGTGGGCGGACACCAAAAACACCGAAATACCC ATACGACGTCCCAGACTACGCCTACCCATATGAT GTCCCGGATTACGCTTACCCATACGATGTTCCAG ATTACGCTTAATATAAATACTTCATTTATTCCATAT GTTTATATTTCA | TG4584, TG4894, TG4969, TG4968, TG4828, TG4896, TG4897, TG4951, TG4953, TG4965 |

**Table S3.** MultiBac expression vectors for LEM-3 production

| Plasmids | MW of LEM-3 protein |
|---|---|
| pFL-His-GST-TEV-GSM-LEM3$^{1-704}$ | 78.2kDa |
| pFL-His-GST-TEV-GSM-LEM3$^{241-704}$ | 52kDa |
| pFL-His-GST-TEV-GSM-LEM3$^{241-704}$-Y530F | 52kDa |
| pFL-His-GST-TEV-GSM-LEM-3$^{241-704}$-R456A | 52kDa |
| pFL-His-GST-TEV-GSM-LEM-3$^{241-704}$-K441A-R456A | 52kDa |
| pFL-His-GST-TEV-GSM-LEM3$^{341-704}$ | 41kDa |
| pFL-His-GST-TEV-GSM-LEM-3$^{485-704}$ | 24kDa |

**Table S4.** Sequences of oligonucleotides for synthetic DNA substrates (Wyatt et al. 2013).

| Oligo | Sequence (5'-3') |
|---|---|
| X0-1 | ACGCTGCCGAATTCTACCAGTGCCTTGCTAGGACATCTTTGCCCACCTGCAGGTTCACCC |
| X0-2 | GGGTGAACCTGCAGGTGGGCAAAGATGTCCATCTGTTGTAATCGTCAAGCTTTATGCCGT |
| X0-3 | ACGGCATAAAGCTTGACGATTACAACAGATCATGGAGCTGTCTAGAGGATCCGACTATCG |
| X0-4 | CGATAGTCGGATCCTCTAGACAGCTCCATGTAGCAAGGCACTGGTAGAATTCGGCAGCG T |
| X26-1 | GCGCTACCAGTGATCACCAATGGATTGCTAGGACATCTTTGCCCACCTGCAGGTTCACCC |
| X26-2 | GGGTGAACCTGCAGGTGGGCAAAGATGTCCTAGCAATCCATTGTCTATGACGTCAAGCTC |
| X26-3 | GAGCTTGACGTCATAGACAATGGATTGCTAGGACATCTTTGCCGTCTTGTCAATATCGGC |
| X26-4 | GCCGATATTGACAAGACGGCAAAGATGTCCTAGCAATCCATTGGTGATCACTGGTAGCGC |
| X0-2.5 | GGGTGAACCTGCAGGTGGGCAAAGATGTCC |
| X0-3.5 | CATGGAGCTGTCTAGAGGATCCGACTATCG |
| X0-1.32 | ACGCTGCCGAATTCTACCAGTGCCTTGCTAGG |
| X0-1.28** | ACATCTTTGCCCACCTGCAGGTTCACCC |

ssDNA= X0-1*
Duplex= X0-1* + X0-4
Immobile Holliday Junction (HJ)-HJ X0-1 = X0-1* + X0-2 + X0-3 + X0-4
Immobile HJ-HJ X0-2 = X0-1 + X0-2* + X0-3 + X0-4
Immobile HJ-HJ X0-3 = X0-1 + X0-2 + X0-3* + X0-4
Immobile HJ-HJ X0-4 = X0-1 + X0-2 + X0-3 + X0-4*
Mobile HJ-HJ X26 = X26-1* + X26-2 + X26-3 + X26-4
Nicked HJ= X0-1.32 + X0-1.28** + X0-2 + X0-3* + X0-4
Replication Fork (RF)= X0-1* + X0-2.5 + X0-3.5+ X0-4
5'-flap= X0-1*+ X0-2.5 + X0-4
3'-flap=X0-1*+ X0-3.5 + X0-4
* The oligonucleotide is labelled by 5'-Cy5
** This oligonucleotide carries a 5'-phosphate group
